# Chronic oxytocin-driven alternative splicing of CRFR2α induces anxiety

**DOI:** 10.1101/2020.08.19.255844

**Authors:** Julia Winter, Magdalena Meyer, Ilona Berger, Sebastian Peters, Melanie Royer, Marta Bianchi, Simone Stang, Dominik Langgartner, Stefan O. Reber, Kerstin Kuffner, Anna K. Schmidtner, Finn Hartmann, Anna Bludau, Oliver J. Bosch, David A. Slattery, Erwin H. van den Burg, Inga D. Neumann, Benjamin Jurek

## Abstract

Recently, oxytocin (OXT) has generated considerable interest as potential treatment for psychiatric disorders, including general anxiety disorder or autism spectrum disorder. Therefore, knowledge on the involved molecular processes downstream of OXT receptor (OXTR) activation is indispensable. We reveal that alternative splicing of corticotropin releasing factor receptor 2α (CRFR2α) parallels increased anxiety-like behavior following chronic OXT treatment, contrasting the well-known anxiolysis of acute OXT. In detail, chronic OXT shifts the splicing ratio between membrane-bound (mCRFR2α) and soluble CRFR2α (sCRFR2α) in favor of the latter via ERK1/2-MEF2A signaling. Targeted manipulations of *Crfr2α* splicing mimic the effect of chronic OXT, confirming its role in the regulation of anxiety-like behavior. Furthermore, chronic OXT triggers cytoplasmic distribution and extracellular release of sCRFR2α into the cerebrospinal fluid, with sCRFR2α levels positively correlating with anxiety-like behavior. Concluding, the dichotomy between anxiolytic mCRFR2α and anxiogenic sCRFR2α is the basis for the deleterious effects of chronic OXT on anxiety.

**Graphical Abstract:** 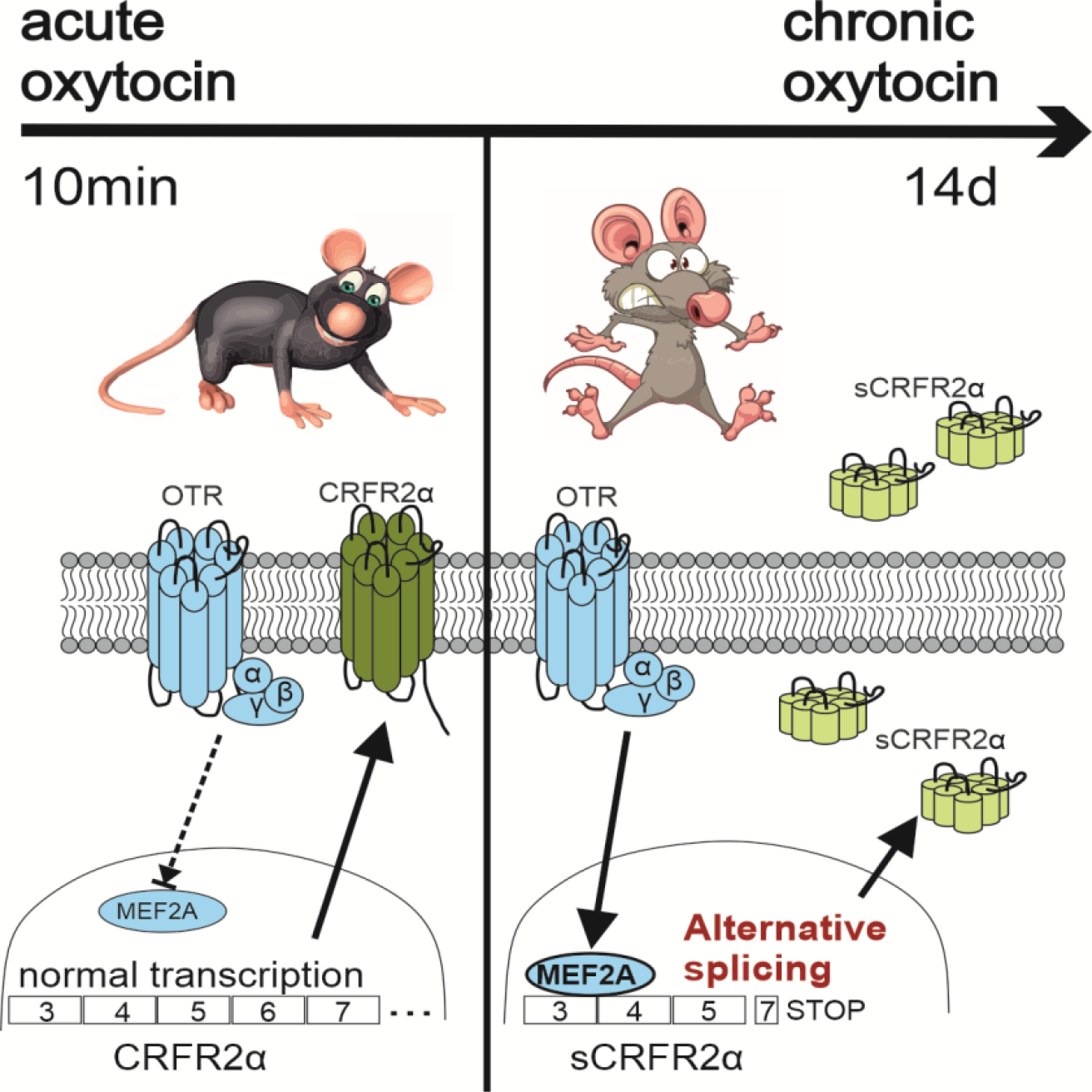

## INTRODUCTION

Neuropeptides and their receptors are established modulators of neuronal activity, shaping multiple behavioral and physiological responses to environmental stimuli. Neuropeptide signaling is therefore an important target for the development of pharmacological treatments of psychopathologies (Hokfelt et al., 2003; Hoyer and Bartfai, 2012). The neuropeptide oxytocin (OXT) has received considerable attention over the last decades due to its acute prosocial and anxiolytic effects, both in humans and animals. Studies in rodents have shown that the prosocial, anxiolytic, and fear-reducing activity of OXT can be induced in the paraventricular nucleus of the hypothalamus (PVN) (Blume et al., 2008; Jurek et al., 2012), central amygdala (Bale et al., 2001; Knobloch et al., 2012; Viviani et al., 2011), lateral septum (Menon et al., 2018; Zoicas et al., 2014), prelimbic and anterior cingulate cortex (Sabihi et al., 2014), and brainstem (Yoshida et al., 2009). Moreover, OXT has been associated with beneficial effects in various physiological and inflammatory processes, such as COVID19 (Soumier & Sirigu 2020), inflammatory pain processing (Eliava et al 2016), and various forms of carcinoma and sarcoma (Reversi et al., 2006).

On a molecular level, upon activation, the G protein-coupled OXT receptor (OXTR) mediates its behavioral effects by multiple intraneuronal signaling cascades (Busnelli and Chini, 2018; Jurek and Neumann, 2018). In brief, the G protein α_q_ subunit is at the basis of activation of protein kinase C (PKC) (Martinetz et al., 2019), whereas the β/γ subunit activates Ca^2+^-influx through transient receptor potential vanilloid type 2 (TRPV2) channels (van den Burg et al., 2015). PKC and Ca^2+^ transactivate the epidermal growth factor receptor (Blume et al., 2008), which is followed by the recruitment of mitogen-activated protein kinase (MAPK) kinase (MEK1/2) signaling (van den Burg et al., 2015) and protein synthesis (Martinetz et al., 2019). Both MEK1/2 signaling and protein synthesis are necessary for the acute anxiolytic effect of OXT in the PVN of male (Blume et al., 2008; Martinetz et al., 2019) and virgin female (Jurek et al., 2012) rats. The anxiolytic activity of OXT in the PVN might be enhanced under mild stress conditions (Blume et al., 2008), indicating an interaction of OXT with central stress regulation. Indeed, OXT has been found to reduce and delay the expression of corticotropin releasing factor (*Crf*) in the hypothalamus (Jurek et al., 2015; Smith et al., 2016), and reciprocal interactions with the anxiolytic (Bale et al., 2000; Deussing and Chen, 2018) transmembrane CRF receptor 2α (CRFR2α) signaling in the PVN and bed nucleus of the stria terminalis have been described (Dabrowska et al., 2011; Janecek and Dabrowska, 2019).

Although OXT is generally considered to be an anxiolytic and pro-social neuropeptide (Donaldson and Young, 2008; Jurek and Neumann, 2018; Neumann and Slattery, 2016), we and others have found adverse effects on anxiety (Peters et al., 2014), fear (Guzman et al., 2013), and social behaviors (Bales et al., 2013; Huang et al., 2014; Pagani et al., 2020), especially after chronic OXT application or virally enhanced OXT signaling. For example, we have recently observed a dose-dependent anxiogenic effect in mice following 15 days of continuous intracerebroventricular (i.c.v.) infusion of OXT (10 ng/h/0.25 µl) using osmotic minipumps (Peters et al., 2014). Moreover, viral vector-induced overexpression of the OXTR in the lateral septum enhanced contextual fear in socially defeated mice (Guzman et al., 2013). However, the molecular mechanisms underlying the adverse effects of chronic or prolonged OXT on anxiety, fear and social behavior, in contrast to its acute effects, have been mostly disregarded. Nevertheless, these conflicting effects have to be carefully considered before OXT can be used as a treatment option for psychiatric disorders (Jurek and Meyer, 2020; Meyer-Lindenberg et al., 2011; Neumann and Slattery, 2016; Winter and Jurek, 2019). Therefore, a deeper understanding of the underlying molecular mechanisms of these possible side effects is indispensable.

To better understand the apparent anxiogenic effects of chronic OXT, we have administered OXT i.c.v. over 14 days in adult rats and found this to enhance anxiety in both males and females. Uniquely in males, increased anxiety is brought about by a novel OXTR-mediated intracellular signaling pathway in the PVN. This pathway includes the MAPK-controlled phosphorylation of the transcription factor myocyte enhancer factor 2A (MEF2A) and subsequent alternative splicing of CRFR2α to its soluble form (sCRFR2α, Chen et al., 2005), which appeared to have a strong anxiogenic effect. Taken together, our data provide a mechanistic explanation of the anxiogenic effect of OXT, and illustrate the intimate relationship between the OXT and CRF systems. Consequently, they challenge the concept of treating anxiety disorders with repetitive or chronic OXT application (Alaerts et al., 2019; Hurlemann, 2017; Neumann and Slattery, 2016; Spengler et al., 2017; Tachibana et al., 2013), and forward sCRFR2α as a novel target for the development of anxiolytics.

## RESULTS

### Chronic OXT infusion dose-dependently induces an anxiogenic phenotype in male and female rats

To assess whether chronic OXT influences anxiety-like behavior, we infused OXT (1 ng/h or 10 ng/hour, 0.25 µl/h) i.c.v. with the aid of osmotic minipumps (Alzet, model 1002) in adult male and female rats over 14 days. The higher dose of 10 ng/h has been reported to modulate anxiety-like behavior in mice (Peters et al., 2014) and rats (Slattery and Neumann, 2010), to reduce intracerebral OXTR expression (Huang et al., 2014; Peters et al., 2014), and has been predicted to saturate brain OXTR over the entire course of the chronic OXT treatment (Chini et al., 2017). Following chronic OXT delivery, rats were tested for anxiety-like behavior in the light-dark box (LDB). In previous studies, mild stress reinforced the acute anxiolytic effect of OXT (Blume et al., 2008); therefore, 24 h prior to LDB testing, we exposed all vehicle (VEH; Ringer’s solution) or OXT-infused rats to a mild stressor by placing the animals on an elevated platform for 5 min (Fig. 1A). When compared to VEH controls, male rats treated with the high, but not low, dose of OXT spent less time in the light box demonstrating an anxiogenic effect of chronic OXT (Fig. 1B, 1C). Remarkably, the observed increase in anxiety-like behavior contrasted with the anxiolytic effect seen after acute OXT infusion into the PVN under otherwise identical experimental conditions (Fig. 1D, E), and with the absence of any effect on anxiety-like behavior, when acutely infused i.c.v. (Fig. S1A, Slattery and Neumann, 2010). The anxiogenic phenotype was reversed five days after termination of chronic OXT infusion demonstrating that OXT-induced anxiogenesis is transient in nature (Fig. S1B). In females, the lower dose increased anxiety-like behavior, while the higher dose, anxiogenic in males, was without significant effects in the LDB (Fig. S1C). This might indicate a higher sensitivity to chronic OT in females, or that intracellular signaling downstream of the OXTR is different between sexes. Importantly, OXT-induced anxiogenesis was not accompanied by changes in locomotor activity, either in males or in females (Table S1). Thus, chronic i.c.v. infusion of OXT dose-dependently increases anxiety in both males and female rats, which reverses to pre-drug anxiety levels within five days after OXT delivery had ended.

**Figure 1.**
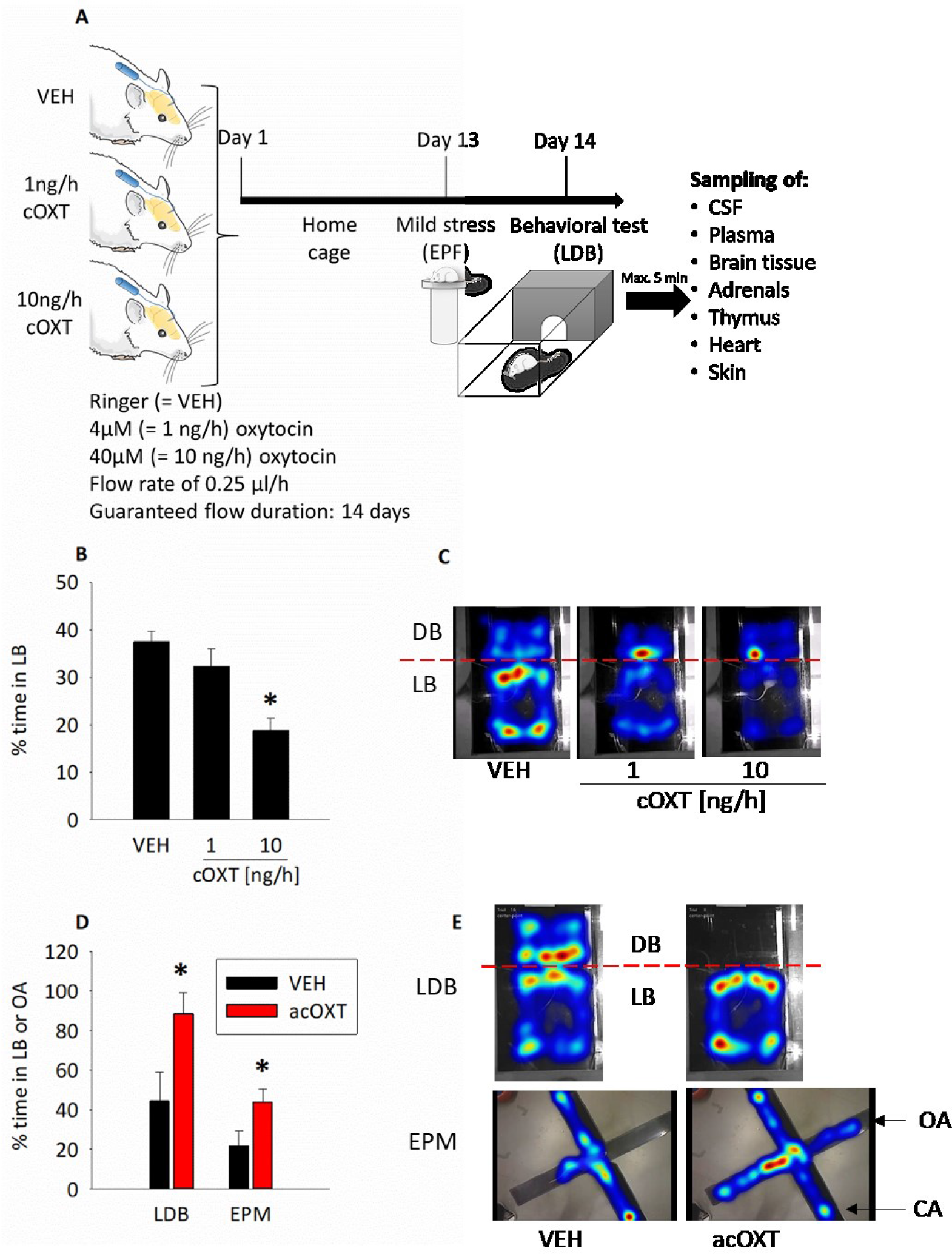
Chronic i.c.v. OXT infusion increases anxiety in male rats. (A) Schematic of the chronic OXT (cOXT) infusion paradigm. Osmotic minipumps (flow rate 0.25 µl/h, infusion duration 14 days) were filled with 4 or 40 µM OXT, corresponding to 1 and 10 ng/h of OXT release rates, respectively, and connected to the lateral ventricle of rats. Day 1 marks the start of infusions. After 13 days, rats were mildly stressed by exposure to the elevated platform (EPF) for 5 min, and tested in the light-dark box (LDB) on day 14. Brain samples were collected immediately after LDB had ended. (B) cOXT decreases time spent in the light box dose-dependently, preceded by EPF. Shown is the percentage of time spent in the lit box (LB). Data are represented as mean + SEM. F_(2;50)_ = 10.131; p < 0.001; Holm-Sidak * p < 0.001 vs VEH; n_(VEH)_ = 15, n_(1ng/h, 10ng/h)_ = 18. (C) Representative heat maps of rat location in the LDB. Treatments as described in (B). (D) Anxiolytic effect of acute bilateral OXT (acOXT; 20 µM, 0.01 nmol/0.5 µl per side) infusion into the paraventricular nucleus (PVN). Male rats were tested in the LDB or elevated plusmaze (EPM) 10 min after infusion, and 24 h after EPF exposure. Shown is the percentage of time spent in the LB or open arm (OA). Data are represented as mean + SEM. One-tailed Student’s *t*-test, p < 0.01, df=6; n_(VEH)_ = 4, n_(acOXT)_ = 4. (E) Representative heat maps of rat location in the LDB and EPM after intra-PVN VEH or acOXT treatment.

### Chronic OXT infusion does not affect the endogenous OXT and AVP systems, nor the hypothalamo-pituitary-adrenal (HPA) axis

To address the possibility that the anxiogenic effect of chronic OXT infusion is due to downregulation of the brain OXT system including its receptor, as seen in mice and voles (Bales et al., 2013; Huang et al., 2014; Peters et al., 2014), or to dysregulation of the closely-related arginine vasopressin (AVP) system, we quantified the expression of OXT, OXTR and V1aR as well as receptor binding within numerous brain areas including PVN. However, none of these parameters of the endogenous OXT or AVP systems were affected by chronic OXT infusion in male rats (Table S2).

As acute and chronic OXT treatment have been linked to stress responses and hypothalamo-pituitary-adrenal (HPA) axis activation (Jurek et al., 2015; Neumann et al., 2000b; Peters et al., 2014; Windle et al., 2004), we monitored selected stress-related parameters including plasma adrenocorticotropic hormone (ACTH) and corticosterone (CORT), as well as adrenal, thymus and heart weights, but found them to be unaltered by chronic OXT (Table S3). Whereas the absence of effects on stress-related parameters, as measured after the entire OXT infusion period, could suggest that HPA axis activity is not involved in OXT-induced anxiogenic effects, we did notice a necessity for a mild stressor to enhance anxiety (Fig. S1D). Transient stress- or arousal-related factors could therefore still play a permissive role to enhance anxiety-related behavior.

### Chronic OXT treatment recruits the transcription factor MEF2A in the PVN of male rats

OXT is released somatodendritically within the rat PVN in response to stressful and fear-enhancing stimuli (Engelmann et al., 2004; Landgraf and Neumann, 2004; Neumann, 2007). Moreover, local OXT controls anxiety-like behavior by modulating intracellular signaling coupled to the OXTR (Blume et al., 2008; Jurek et al., 2015; Jurek et al., 2012; van den Burg et al., 2015). Therefore, we aimed to resolve the biochemical mechanism in the PVN that may underlie chronic OXT-induced anxiogenesis. In PVN protein extracts, we observed increased phosphorylation, reflecting activation, of the MAP kinases MEK1/2 (at 1 and 10 ng/h) and ERK1/2 (at 10 ng/h) after 14 days of OXT infusion in males, whereas no such activation was found in females, (Figs. 2A, B, S2A, B).

**Figure 2.**
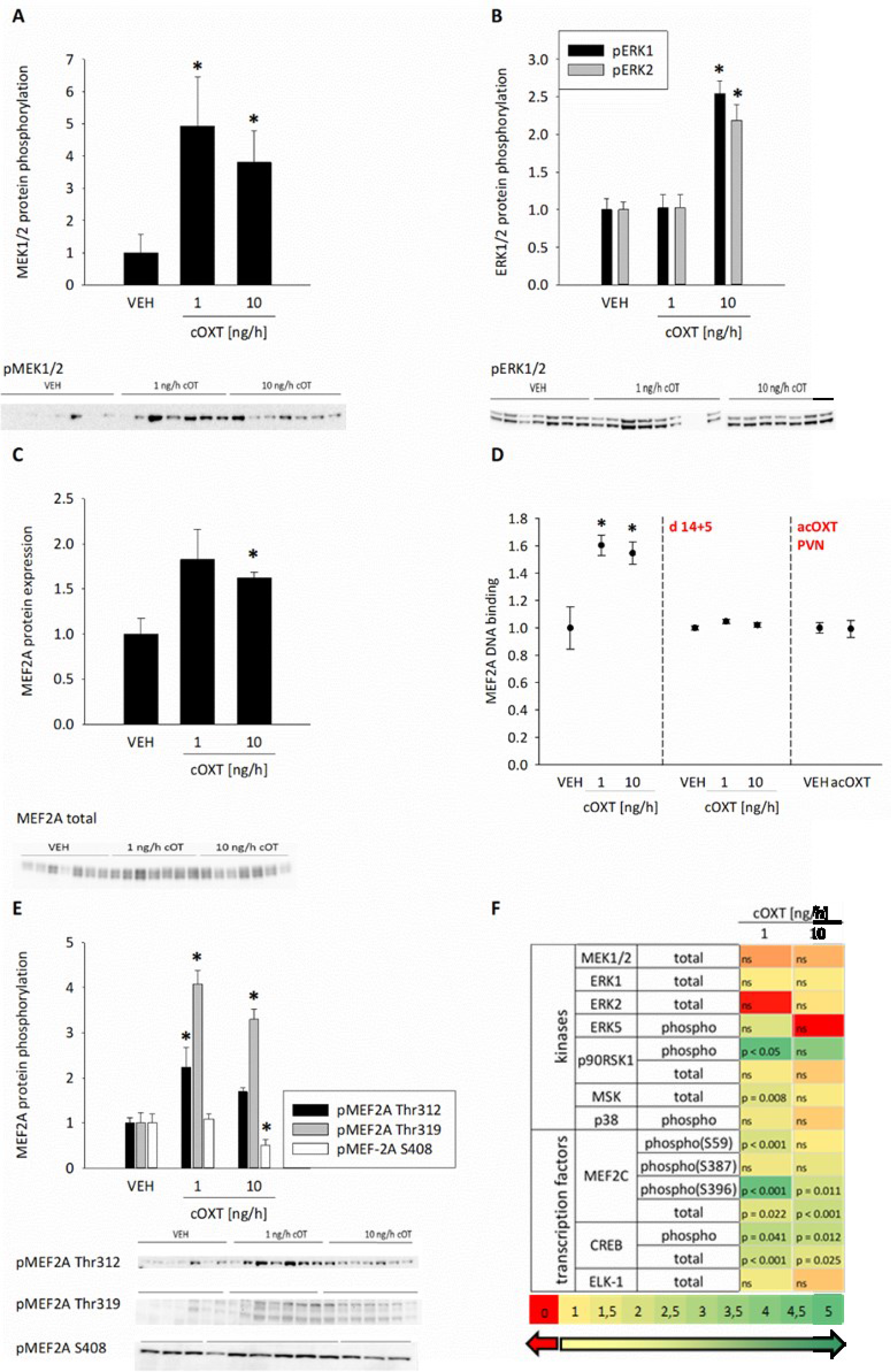
Recruitment of MAPK and MEF2A by chronic OXT infusion in the PVN of male rats. (A) cOXT increases MEK1/2 phosphorylation. Data are represented as mean +SEM. Kruskal-Wallis H = 8.396; p = 0.015; Dunn’s Method * p < 0.05 vs VEH; n_(VEH; 1ng/h; 10ng/h)_ = 7 (B) ERK1/2 phosphorylation increase in the 10 ng/h cOXT group; n_(VEH; 1ng/h; 10ng/h)_ = 7; pERK1: F_(2;20)_ = 9.672; p = 0.001; Holm-Sidak * p = 0.001 VEH, 1 ng/h vs 10 ng/h; pERK2: Kruskal-Wallis H = 10.293 p = 0.006; Tukey Test VEH vs 10 ng/h * p < 0.05, 1 ng/h vs 10 ng/h * p < 0.05. (C) MEF2A total protein expression increases in PVN lysates of 10 ng/h cOXT-treated and mildly stressed male rats. Kruskal-Wallis H = 7.156, p = 0.028; Dunn’s Method *p < 0.05 VEH versus 10ng/h; n_(VEH)_ = 6, n_(1ng/h)_ = 6, n_(10ng/h)_ = 7. (D) MEF2A DNA binding activity increases in PVN lysates after 14d chronic OXT and mild stress. Values are normalized to VEH. F_(2;39)_ = 6.732, p = 0.001, Holm-Sidak * p < 0.05 vs VEH; n_(VEH)_ = 7, n_(1 ng/h)_ = 7, n_(10 ng/h)_ = 16. In contrast to 14 days of cOXT treatment, MEF2A binding activity returned to basal, 5 days after the infusion stopped (cOXT d14+5). Data are represented as the fold changes in DNA-binding activity. F_(2,14)_ = 2.316, p = 0.141; n_(VEH)_ = 5, n_(1ng cOXT)_ = 5, n_(10ng cOXT)_ = 5. No effects of acute intra-PVN OXT infusions have been observed on local MEF2A binding activity. Data are represented as fold changes in DNA binding activity + SEM. t = 0.0938; two-tailed p-value = 0,927, n_(VEH)_ = 6, n_(OXT)_ = 7. (E) Phosphorylation of Thr312 and Thr319 within MEF2A increase, and that of S408 decreases after cOXT and mild stress, indicating increased activity. pMEF2A Thr312: Kruskal-Wallis H = 9,106, p = 0.011; Tukey Test VEH vs 1 ng/h * p < 0.05. pMEF2A Thr319: F(2;21) = 8.887; p = 0.006; Holm-Sidak * p = 0.002 VEH versus 1 ng/h and 10 ng/h. pMEF2A S408: F(2;23)=4.614; p = 0.022; Holm-Sidak * p = 0.033 VEH versus 10 ng/h cOXT. (F) Heat map of expression and phosphorylation levels of second messenger kinases and transcription factors downstream of the OXTR as analyzed by Western blot. Fold change below 1 (red) indicates downregulation, fold change of 1 (yellow) indicates no change, fold change bigger than 1 indicates upregulation (green). p-values vs VEH included; n.s. = not significant.

As we have previously demonstrated that OXT for 12 h recruits the transcription factor MEF2A in hypothalamic neurons in a MAPK-dependent manner (Meyer et al., 2018), we assessed whether chronic OXT infusion would lead to MEF2A activation. We found that chronic OXT treatment (10 ng/h) resulted in elevated total protein levels of MEF2A (Fig. 2C), and this was indeed accompanied by increased MEF2A activity as indicated by increased MEF2A DNA binding in PVN tissue lysates of OXT-treated male, but not female rats (Fig. 2D, S2C). Phosphorylation of the transcription activation sites Thr312 and Thr319 specifically reflected elevated MEF2A activity, whereas the MEF2A inhibitory site Ser408 was dephosphorylated following chronic OXT (Fig. 2E). Taken together, these results indicate a strong activation of MEF2A in the PVN of male rats that had been treated with OXT for 14 days.

Remarkably, OXTR-coupled signaling underlying the acute anxiolytic effect in the PVN does not involve the activation of MEF2A (Fig. 2E), and an acute i.c.v. OXT infusion failed to stimulate MEF2A activity (Fig. S2D), demonstrating the specificity of MEF2A recruitment following chronic OXT administration. Furthermore, MEF2A DNA binding capacity was similar to that seen in VEH-treated rats five days after i.c.v. chronic OXT infusion had ended and the anxiogenic effect disappeared (Fig. 2E). The upregulation of MEF2A activity and expression in the PVN of chronically OXT-infused male rats appeared to be highly specific, as none of the other related kinases tested, including p38, p90rsk, and MSK, showed any significant changes in phosphorylation (Fig. 2F). The upregulation of MEF2A was also brain region-specific, as neither the hippocampus, prefrontal cortex, amygdala, raphe nuclei, nor septum revealed increased MEF2A/B/C expression following chronic OXT treatment (Table S2 and Fig. S2E). In addition, the phosphorylation pattern of the closely related MEF2C, strongly expressed in the PVN and linked to OXTR signaling (Devost et al., 2008), revealed no change at Ser59, the regulatory site inducing DNA binding (Fig. 2F). Consequently, DNA binding of MEF2C remained unaffected by the behaviorally relevant 10 ng/h OXT treatment (Fig. S2F).

In conclusion, chronic central infusion of high OXT results in MEF2A activation in the PVN of male rats, correlating with increased anxiety-related behavior. Recruitment of MEF2A does not underlie the anxiogenic effect induced by infusion of the low dose of OXT in virgin female rats. The intracellular mechanism causing anxiogenesis in females is distinct from that in males and needs further investigation.

### MEF2A shifts the expression of membrane-bound to soluble CRFR2α in the PVN

The demonstration of the co-occurrence of MEF2A activation and high anxiety-like behavior in male rats after chronic OXT treatment suggests that MEF2A controls the expression of genes that could play a role in anxiogenesis. To identify these, we made use of a custom-designed PCR array that targeted genes in the PVN that 1) are known to be involved in processes related to MEF2A function, such as synaptic connectivity or neuronal plasticity, 2) contain one or more MEF2A binding sequences, and 3) have been associated with stress- or anxiety-like behaviors. After high chronic OXT treatment, we found distinct alterations in gene expression of closely related genes, of which those of membrane-bound and soluble CRFR2α were particularly striking (Fig. 3A). As previous studies have shown interactions between OXT and CRF signaling (Jurek et al., 2015; Li et al., 2016), and as CRF controls stress and fear / anxiety (Asok et al., 2018; Bale et al., 2000; Bale et al., 2002b; Radulovic et al., 1999), we decided to explore a possible role of the CRFR2α in anxiogenesis further.

**Figure 3.**
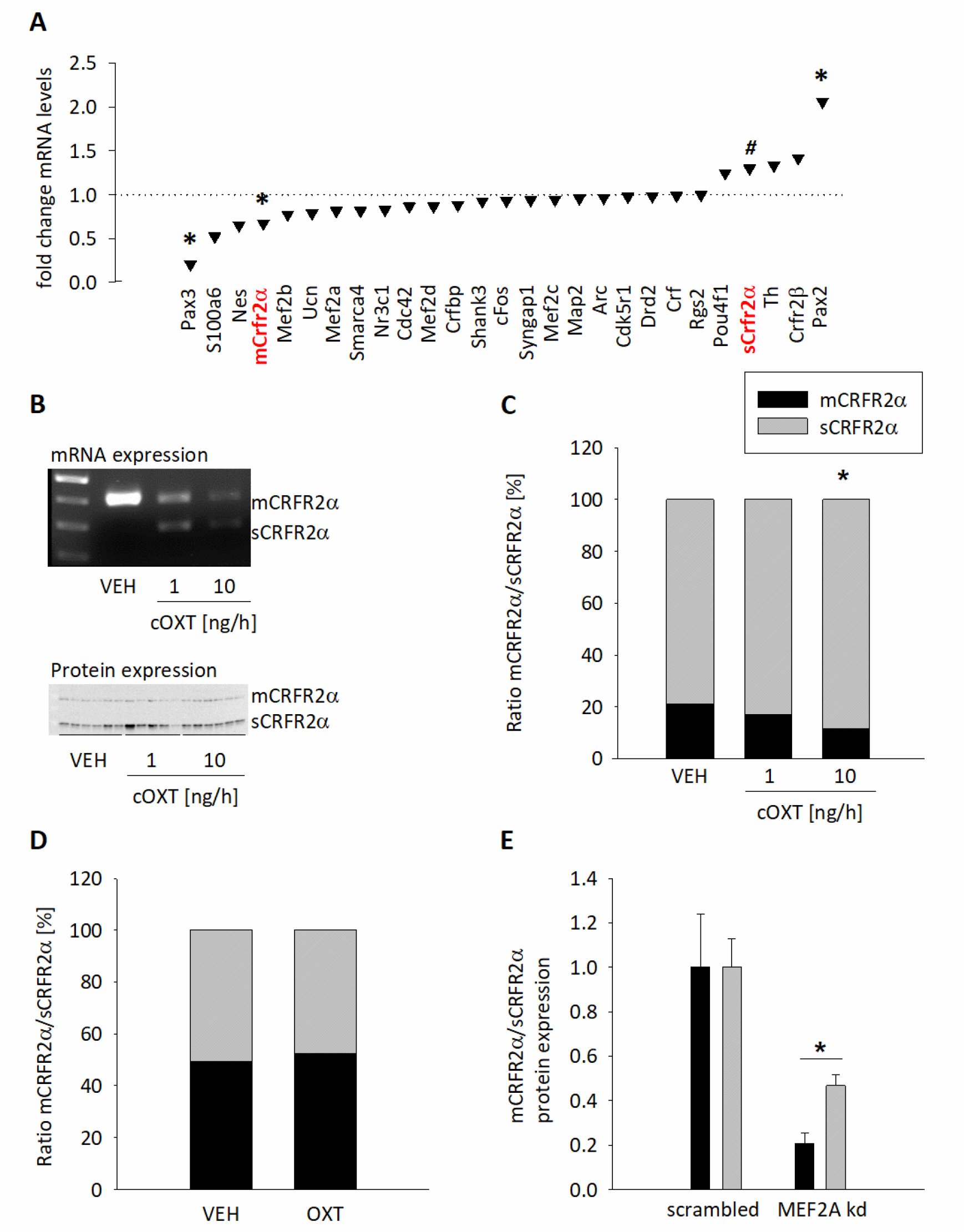
MEF2A-dependent regulation of OXT receptor target genes. (A) PCR array: Expression of MEF2A-regulated candidate genes after 10 ng/h cOXT treatment and mild stress. Data calculated by ΔΔCT and presented as fold change vs. VEH. Fold changes < 1, upregulated mRNA expression, fold changes > 1, downregulated mRNA expression. n_(VEH)_ = 5, n_(10ng/h cOXT)_ = 6. Pax3 * p = 0.021, Crfr2α * p = 0.02, sCrfr2α # p = 0.07, Pax2 * p = 0.05. (B) Upper panel: Representative example of 3 male rat PVN cDNA samples showing mCRFR2α (upper band, 400 bp) in VEH-treated animals, and mCRFR2α / sCRFR2α (lower band, 300 bp) expression in the cOXT treatment groups. Lower panel: Western Blot of mCRFR2α (approx. 50 kDa) and sCRFR2α (approx. 30 kDa) levels after VEH, 1 or 10 ng/h cOXT infusion and mild stress, using a pan-CRFR2 antibody. (C) Ratio of mCRFR2α and sCRFR2α protein expression after cOXT and mild stress in male rats, shifting in the favor of sCRFR2α following treatment. Data are represented as mean + SEM. F_(2;15)_ = 5.311, p = 0.021; Holm-Sidak * p = 0.019 vs VEH; n_(VEH)_ = 6, n_(1 ng/h)_ = 5, n_(10 ng/h)_ = 6. (D) Treatment with acOXT had no effects on the ratio of mCRFR2α/sCRFR2α protein expression in % in the PVN of male rats. Data are represented as mean + SEM. t = 0.784 with 11 degrees of freedom, two-tailed p-value = 0.450; n_(VEH)_ = 6, n_(OXT)_ = 7. (E) siRNA-mediated knockdown of MEF2A in H32 cells incubated with 100 nM OXT decreased mCRFR2α and sCRFR2α protein levels, indicating a central role for MEF2A in the transcription of the CRFR2 gene. Data represented as mean + SEM. mCRFR2α: Mann-Whitney < 0.001, * p = 0.002; sCRFR2α: t = 2.574 with 10 degrees of freedom; one-tailed * p-value = 0.0138, both vs. respective scr (scrambled) control; n_(scr)_ = 6, n_(MEF2A kd)_ = 6.

The expression of membrane-bound CRFR2α (mCRFR2α; Fig. 3A) was significantly reduced in 10 ng/h OXT-treated rats, whereas that of the alternative splice variant, i.e. soluble sCRFR2α, was increased. The alternative splicing was confirmed by endpoint PCR demonstrating that sCRFR2a expression, which was below the detection limit in VEH controls, is effectively induced after chronic OXT infusion (Fig. 3B upper panel). At the protein level, OXT-treated animals expressed significantly more sCRFR2α (88.3%) in proportion to the mCRFR2α (11.7%; Fig. 3B lower panel and 3C). The increased sCRFR2α over mCRFR2α protein ratio was exclusively mediated by increased sCRFR2α, as mCRFR2α remained stable (data not shown). Thus, chronic OXT likely promotes alternative splicing of CRFR2α to its soluble form in males (Fig. 3B, C), but not in females (Fig. S3A).

As in silico analysis revealed that the coding sequence of the *Crfr2α* gene harbors seven potential MEF2A binding sites, with two being predicted within Exon 6 (base pairs 791-800 and 908-917; Fig. 5B), we hypothesized that MEF2A drives alternative splicing of CRFR2α directly, and elevated sCRFR2α levels mediate the anxiogenic activity of chronic OXT. To test this, we performed chromatin immunoprecipitation (ChIP; Fig. S3B, schematic overview) with rat hypothalamic H32 cells (Mugele et al., 1993), which express MEF2A, OXTR and CRFR2α (Fig. S3C, S3D). We found MEF2A binding to the CRFR2α gene especially within Exons 2 (base pairs 390-399) and 6 (base pairs 791-800 (Fig. 5B). Furthermore, selective MEF2A knockdown by means of siRNA strongly reduced both mCRFR2α and sCRFR2α expression in these cells, highlighting the necessity of MEF2A to induce CRFR2α gene transcription (via binding to Exon 2) and to promote alternative splicing (via binding to Exon 6; Fig. 3E and 5B). In confirmation of the presence of the OXTR-MEF2A-sCRFR2α cascade within OXTR-expressing PVN neurons, we found that OXTR (as revealed by a Venus signal) and sCRFR2α are partially co-expressed in the PVN of OXTR reporter mice (Yoshida et al., 2009). Co-localization between vasopressin 1A and 1B receptors on the one hand, and sCRFR2α on the other hand, appeared to be quite rare; thus, any contribution of the vasopressin system to the observed biochemical and behavioral effects is expected to be negligible (Fig. S3H and J).

### Chronic OXT promotes the release of sCRFR2α

After establishing that chronic OXT treatment alters MEF2A activity and, subsequently, promotes alternative splicing of the *Crfr*2α gene and sCRFR2α protein synthesis, we next sought to determine whether prolonged OXT application (i.e. 24 h) in hypothalamic cell culture can stimulate the release of the sCRFR2α protein into the extracellular medium. To this end, we integrated a spliceable tag (HiBiT) into Exon 6 of CRFR2α by means of CRISPR-Cas9 technology in H32 cells. This tag was visualized by adding a “LgBiT” substrate, the binding protein of the HiBiT-tag, to the cell culture medium (Fig. 4A). The luminescent signal that forms following binding LgBiT to HiBiT can be measured in real time and detects membrane-bound mCRFR2α. We found that luminescence intensity dropped in cells that had been exposed to 100 nM OXT, when compared with VEH-treated cells, indicating reduced membrane incorporation of mCRFR2α following chronic OXT exposition (Fig. 4B). In parallel, sCRFR2α content increased tenfold in the cell culture medium as assessed by Dot Blot analyses using a specific sCRFR2α antibody (Chen et al., 2005), Fig. 4C, S3D). This indicates, for the first time, that OXT induced the release of CRFR2α from hypothalamic H32 cells. In addition, immunohistochemical stainings of primary hypothalamic cells revealed punctate sCRFR2A immunoreactivity in neurons, reflecting vesicular localization (Fig. 4D).

**Figure 4.**
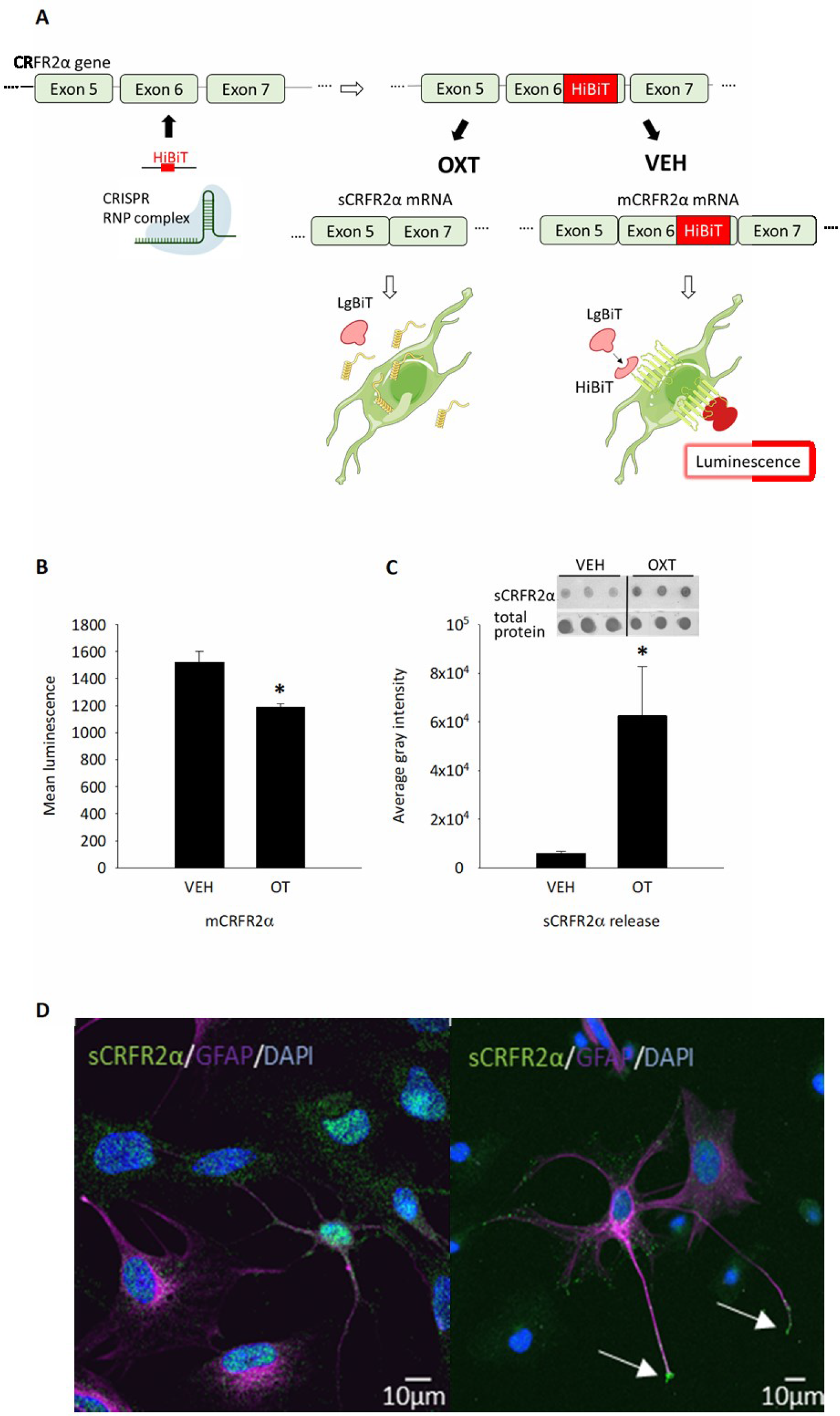
Chronic OXT promotes the release of sCRFR2α *in vitro*. (A) Schematic representation of HiBiT-mediated signaling in H32 cells stimulated with VEH or OXT for 24 h. (B) Assessment of mCrfr2α membrane expression by tagging the transmembrane domain encoded in Exon 6 with a HiBiT-tag. Membrane expression is indicated by luminescence caused by extracellular HiBiT-LgBiT interaction. Stimulation for 24 h with 100 nM OXT decreased the luminescent signal, indicating alternative splicing of Exon 6 and, therefore, reduced membrane expression of CRFR2α. Data presented as absolute values of mean luminescence +SEM. Mann-Whitney Rank Sum Test, p ≤ 0.001, n_(VEH)_ = 20, n_(OXT)_ = 12. (C) Cell culture supernatants taken from HiBiT-expressing cells (as shown in B) reveal sCRFR2α release into the cell culture medium, which was 10-fold higher after stimulation with 100 nM OXT over 24 h. Data shown as average gray intensity relative to total protein Ponceau red staining. Mann-Whitney Rank Sum Test, p = 0.029; n_(VEH/OXT)_ = 4. Right panel: Representative Dot Blot of triplet sCRFR2α staining and Ponceau red loading control. (D) Left panel: Representative images from rat hypothalamic mixed primary cultures stained for DAPI (blue), sCRFR2α (green), and GFAP (magenta) reveals cytoplasmic distribution of sCRFR2a in neuronal cells (GFAP negative) and astrocytes (GFAP positive). Right panel: Accumulation of sCRFR2α in end-boutons of astrocytic processes (white arrows) might indicate releasable pools.

### Altered sCRFR2α / mCRFR2α ratio increases anxiety in male rats

To determine whether sCRFR2α is secreted into the CSF in living rats, we sampled CSF from the cisterna magna of male rats, and assessed the presence or absence of sCRFR2α. We found detectable amounts of sCRFR2α in the extracellular fluid (Fig. 5A). No sCRFR2α could be detected in the cerebellum (CB) negative control (Chen et al., 2005), confirming the specificity of the dot blot analysis. Following this, we specifically knocked down the sCRFR2α splice variant in the PVN of living rats using locked nucleic acid antisense oligonucleotides, so-called “GapmeRs” (Migawa et al., 2019; Xing et al., 2014) and assessed whether the availability of extracellular sCRFR2α affects anxiety. The GapmeRs were designed to bind the Exon 5-7 boundary, which is exclusively found in the mRNA of the soluble form of CRFR2α, so that specific knockdown of sCRFR2a was achieved (Fig. 5B). Fluorescent labeling of the infused GapmeRs revealed mostly intra-PVN cell body staining, with some diffuse signal scattered in the adjacent, mostly dorsal hypothalamus, potentially representing cellular projections originating from the PVN (Fig. S4A). Importantly, acute infusion of GapmeR into the PVN caused a significant reduction of sCRFR2α in the CSF after 7 days (Fig. 5C), accompanied by a reduction in anxiety-like behavior in two independent behavioral tests (Fig. 5D). Thus, the decrease in anxiety-like behavior in GapmeR-infused rats was reflected both by an increased time spent in the light box of the LDB, as well as an increased exploration of the center area of the open field arena (Fig. 5E and F). In a complementary approach, we upregulated sCRFR2α levels by designing another set of antisense oligonucleotides (referred to as “target site blockers” (TSB)) to provoke alternative splicing of the CRFR2α mRNA exclusively to its soluble form. This was achieved by targeting TSB to the endogenous splice sites up- and downstream of Exon 6, thus enforcing the exclusion of Exon 6 from the pre-mRNA and inducing a frame shift to form a premature termination codon within Exon 7 (Fig. 5B). TSB infusion bilaterally into the PVN resulted in increased anxiety-related behavior of male rats reflected by a reduction of the time spent in the light compartment of the LDB from 50% to 25% (Fig. 5G, 5H). Taken together, these results indicate that a high sCRFR2α / low mCRFR2α ratio in the PVN of male rats is associated with an anxiogenic phenotype. Given the observed increase in *sCrfr2α* gene expression in chronically OXT-infused rats (Fig. 3B), such altered ratio is likely to underlie the anxiogenic effect of chronic OXT treatment. Of note, this effect appears to be specific for anxiety, as social preference behavior, known to be promoted by OXT (Lukas et al., 2011; Menon et al., 2018), was not affected by GapmeRs or TSB (Fig. S4B).

**Figure 5.**
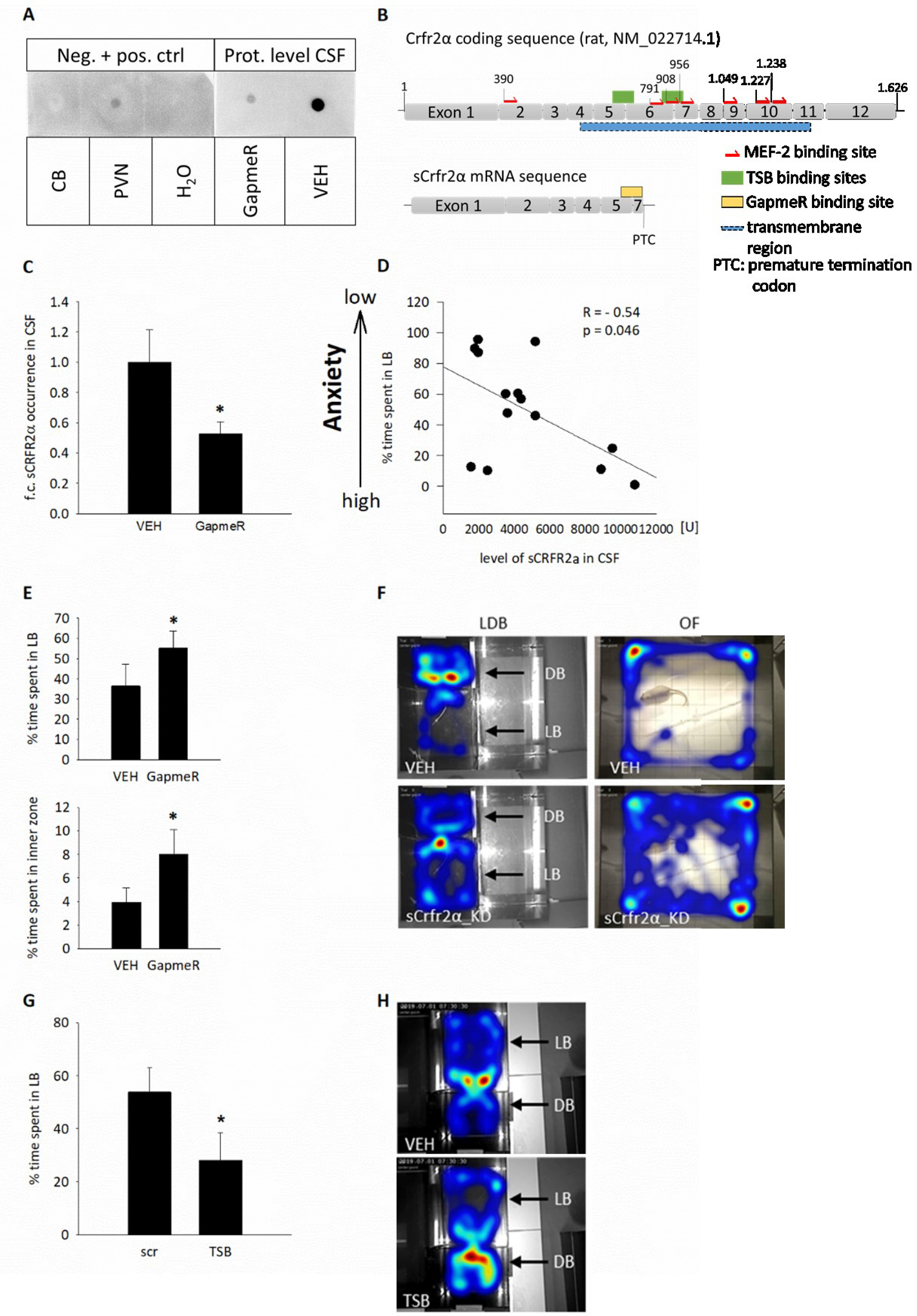
Increased sCRFR2α release underlies the anxiogenic effect of chronic OXT in combination with mild stress. (A) Representative picture of Dot Blot analysis of CSF (VEH and GapmeR) and control tissue protein lysates (cerebellum = CB as negative control, PVN as positive control, H_2_O). (B) Schematic representation of the rat Crfr2α gene, the transmembrane domain is labeled in blue, including Exon 4 to Exon 11. Target site blocker (TSB) binding sites are labeled green. Multiple MEF2 binding sites are indicated as red arrows. Exon 5/7 boundary GapmeR binding site in the sCRFR2 mRNA is indicated in yellow. (C) GapmeR infusion reduced sCRFR2α expression by approximately 50% as assessed by Dot Blot in CSF samples. Data are represented as fold changes in protein expression + SEM. One-tailed Student’s t-test, t = 1.828, df = 12, p = 0.046; n_(scr)_ =8, n_(GapmeR)_ = 6. (D) CSF sCRFR2α levels and anxiety-like behavior (% time spent in LB) correlate negatively (R = −0.54, ANOVA F_(1;13)_ = 4.946, p = 0.046), suggesting that sCRFR2α is anxiogenic. (E) GapmeRs increased time spent in LB and inner zone of open field (OF), indicating anxiolysis. Anxiety-like behavior was determined 7 (LDB) or 8 days (OF) after GapmeR infusion. Data are represented as mean percentage of time spent in the LB + SEM (one-tailed t-test * p = 0.0486), and the mean percentage of time in the inner zone (IZ) of the OF (Mann-Whitney U = 16.000, * p = 0.037) + SEM; n_(scr)_ = 8, n_(GapmeR)_ = 6. (F) Representative heat maps showing the distribution of rat presence in the light and dark compartment of the LDB or inner and outer zone in the OF test. (G) Target site blocker (TSB) treatment enhances anxiety-like behavior of male rats in the LDB 7 days after infusion, demonstrating an anxiogenic effect of sCRFR2α. Data are represented as mean percentage of time spent in the LB + SEM. t = 1.843; one-tailed t-test * p = 0.0409; n_(scr)_ = 10, n_(TSB)_ = 10. (H) Representative heat map of the distribution of rat presence in the LDB after VEH or TSB treatment.

Finally, we assessed whether agonism and antagonism of mCRFR2α in the PVN influences anxiety-like behavior. Infusion of the mCRFR2α-specific agonist stresscopin, which exhibits a 10-fold higher affinity for mCRFR2 than for sCRFR2 (Chen et al., 2005; Deussing and Chen, 2018), increased the time spent in the light box of the LDB (Fig. S4C). In contrast, the mCRFR2α-specific antagonist antisauvagine-30, which binds transmembrane regions of the mCRFR2 and, therefore, not the sCRFR2α (Liaw et al., 1997; Perrin et al., 2003), reduced the time spent in the light box, indicating that functional mCRFR2α signaling in the PVN is necessary for low anxiety levels (Fig. S4C). Therefore, as chronic OXT shifts the balance between sCRFR2α and mCRFR2α in the favor of anxiogenic sCRFR2α, the novel OXTR - ERK1/2 -MEF2A - sCRFR2α biochemical pathway that we found may be at the basis of the anxiogenic effect of OXT in male rats.

## DISCUSSION

Although the anxiolytic effect of acute OXT application and underlying molecular mechanisms have been well characterized (Blume et al., 2008; Jurek et al., 2012; Neumann et al., 2000a; Okimoto et al., 2012; Sabihi et al., 2014; van den Burg et al., 2015), only little is known regarding possible adverse effects of chronic OXT treatment. In this context, detailed knowledge on the molecular processes downstream of the OXTR after chronic OXT is indispensable, as OXT has the potential as treatment option for psychopathologies associated with symptoms of anxiety (Andari et al., 2018; Macdonald and Feifel, 2012; Meyer-Lindenberg et al., 2011; Neumann and Slattery, 2016).

In the current study, we revealed that chronic i.c.v. OXT infusion for 14 days increases anxiety-like behavior in male and virgin female rats. We are the first to show that the anxiogenic effect is accompanied by the activation of a yet unknown intracellular signaling pathway that involves increased transcriptional activity of MEF2A, and subsequent alternative splicing of the CRFR2α into its soluble and anxiogenic form in male, but not female rats. The distinct role of CRFR2α in anxiety regulation could be confirmed by targeted manipulations of *Crfr2α* splicing: Whereas knock down of sCRFR2α within the PVN using GapmeRs reduced anxiety, local upregulation of alternative splicing of CRFR2α exclusively to its soluble form using target site blockers resulted in anxiogenesis, thus mimicking the effect of chronic OXT. We could further show that chronic OXT induced extracellular release of sCRFR2α, which can be quantified in the CSF, positively correlates with anxiety-like behavior. Although our results challenge the utility of chronic or repetitive OXT application as treatment option for anxiety-related disorders, especially in male patients, we could identify sCRFR2α as a novel target for diagnosis and pharmacological intervention of these disorders.

The identified anxiogenic effect of chronic i.c.v. OXT is in contrast to the established anxiolytic effect of an acute OXT bolus. However, anxiolysis in rodents has only been described after OXT infusion directly into distinct brain regions (PVN, central amygdala, septum, prefrontal cortex or brainstem; for review see Jurek and Neumann, 2018). Indeed, we did not observe any effect on anxiety-like behavior 10 min after i.c.v. infusion of OXT, which was paralleled by the absence of MEF2A activation in the PVN. Nevertheless, our data indicate that the PVN may be critically involved, since activation of MEF2A after chronic OXT was restricted to the PVN. However, we cannot exclude the contribution of altered inputs to the PVN to the observed biochemical changes.

The expression of the CRFR2α is restricted to the brain, with few exceptions in the periphery like dermal or heart tissue (Lovenberg et al., 1995; Van Pett et al., 2000). Its shorter splice variant sCRFR2α is unable to incorporate into the cellular membrane due to the lack of the transmembrane domain and, therefore, displays a random cytoplasmic localization, from where it can be released into the extracellular space (Chen et al., 2005; Evans and Seasholtz, 2009). Using an antibody that detects both the full-length mCRFR2α and the splice variant sCRFR2α, we revealed that chronic OXT promoted sCRFR2α expression, whereas mCRFR2α levels remained unchanged, resulting in a shift of the sCRFR2α to mCRFR2α ratio. On transcriptional level, even a decrease in *mCrfr2α* mRNA could be detected, as both the full-length receptor and the splice variant derive from the same mRNA. Indeed, regulation of full-length mRNA availability is discussed to be one of the main functions of alternative splicing of the CRFR2α (Evans and Seasholtz, 2009; Markovic and Grammatopoulos, 2009). Also, mRNA processing and alternative splicing are common ways to regulate class B GPCR availability and function (Deussing and Chen, 2018; Furness et al., 2012). This might account for the effects of chronic OXT on the shifted protein ratio towards sCRFR2α that promoted an anxious phenotype in our study.

We could further identify that activation of the transcription factor MEF2A is involved in the process of alternative splicing, induced by chronic OXT, although, MEF2A as a transcription factor is not intuitively linked to this process. However, recent findings support the hypothesis that transcription and alternative splicing are joint and co-dependent processes, since MEF2A is part of a large complex of transcription and splicing-regulating factors, such as Brg1/Brm, BAF47, BAF170, BAF155, MyoD, and a histone acetyl transferase. These factors contribute to the correct transcription and immediate processing of the transcript (Proudfoot et al., 2002; Zhang et al., 2016). It is, therefore, perfectly feasible to infer a parallel role for MEF2A in the transcription and alternative splicing of the *Crfr2α* gene. Decreased phosphorylation of MEF2A at S408 in the PVN indicates a stimulatory effect of chronic OXT on gene transcription and mRNA processing. This is supported by MEF2 binding sites within the splice-relevant Exon 6 of the *Crfr2α* gene. The switch in signaling specificity after chronic OXT compared to acute OXT occurs feasibly due to variations in duration and strength of the signal (Keyse, 2008; Murphy et al., 2004).

Although not investigated in depth, we observed that exposure to a mild stressor, i.e. elevated platform stress 24 h prior to testing, is required to induce anxiogenesis after chronic OXT treatment. This is similar to what has been reported for the acute anxiolytic effects of OXT in the PVN (Blume et al., 2008). Interestingly, mild stress can promote the incorporation of anxiolytic CRFR2α into the membrane of neurons (Slater et al., 2016; Waselus et al., 2009; Wood et al., 2013) without necessarily activating the HPA axis (Bale et al., 2002a; Bazhan et al., 2013). Thus, chronic OXT might interfere with CRFR2α membrane incorporation by shifting the expression of membrane-bound to anxiogenic soluble CRFR2α.

Virgin female rats appeared to be more sensitive to chronic OXT treatment than males, as already the low, but not high, dose of chronic OXT resulted in anxiogenesis. However, on a molecular level, no changes in mCRFR2α/sCRFR2α ratio could be detected in females, as substantial MEF2A activity was missing. While we have not studied this difference between male and virgin female rats further, sexually dimorphic actions of OXT have been repeatedly described, clearly exemplified during lactation, (Menon et al., 2018; Murgatroyd et al., 2015). In addition, OXT-induced recruitment of CRF binding protein in the prefrontal cortex acts anxiolytic in a sex-specific manner, as CRF binding protein attenuates potentiation of postsynaptic layer 2/3 pyramidal cell activity only in male mice and, thereby, brings about anxiolysis (Li et al., 2016).

Our study has revealed an intimate interaction between OXT and CRF signaling, which controls anxiety-like behavior in male, but not female, rats. CRFR2α-mediated signaling has already been linked to both anxiolysis (Henckens et al., 2017; Issler et al., 2014) and anxiogenesis (Radulovic et al., 1999), which is why we assessed the role of CRFR2α in the PVN in anxiety-like behavior further. The RNA-based approach that we adopted to specifically down-regulate either the membrane-bound or soluble form of CRFR2α showed that mCRFR2α expression is linked to reduced anxiety, whereas alternative splicing and shift towards sCRFR2α results in increased anxiety. In confirmation, pharmacological manipulation of the CRFR2α by infusion of a specific agonist or antagonist of the CRFR2α, bilaterally into the PVN, also resulted in opposite effects on anxiety.

The OXT-induced increase in sCRFR2α, which may act as a scavenger of CRFR2α ligands similar to CRF-BP, indicates that the interactions between OXT and CRFR2α signaling are key in the control of anxiety-like behavior of male rodents. Although shorter in length, sCRFR2α maintains its ability to bind the mCRFR2α ligands CRF, urocortin I and urocortin II with nanomolar affinity (Chen et al., 2005; Reyes et al., 2001; Vaughan et al., 1995), and thus could prevent them from binding to mCRFR2α. Consequently, scavenging mCRFR2α ligands by sCRFR2α could weaken or prevent the anxiolytic signal conveyed by mCRFR2α, and promote anxiety-like behavior. Importantly, we have detected sCRFR2α in CSF proving that sCRFR2α is released into the extracellular space in vivo. The observed correlation between CSF sCRFR2α levels and anxiety-like behavior further supports our hypothesis that the balance between soluble and membrane-bound CRFR2α in the PVN in favor of the splice variant is instrumental in promoting anxiety in male rats. Together, these results demonstrate that a shift from mCRFR2α to sCRFR2α evokes an anxiogenic phenotype in male rats.

In conclusion, our study shows that OXT is able to enhance anxiety-like behavior when infused over a longer period of time, and this involves the activation of the transcription factor MEF2A and subsequent alternative splicing of *Crfr2α* mRNA from the full-length membrane-bound to the anxiogenic soluble form. Also, the results show that intimate interactions between two neuropeptidergic systems, such as OXT and CRF/urocortin, can induce important alterations in behavior. Finally, our study forwards a word of caution for the chronic or repetitive use of OXT for the treatment of psychiatric disorders, especially in men. Instead of broadly manipulating the OXT system, targeting specifically OXT-induced sCRFR2α synthesis and release opens a promising avenue for future treatment options.

## EXPERIMENTAL MODEL AND SUBJECT DETAILS

### *In vivo* animal studies

Adult male and female Wistar rats (Charles River, Germany, 250–300 g at the beginning of the experiment) were housed under standard laboratory conditions in groups of 3 (12 h light-dark cycle, 22 – 24° C, lights on at 06:00, food and water *ad libitum*). All animal experiments were performed between 08:00 – 11:00, in accordance with the Guide for the Care and Use of Laboratory Animals by the National Institutes of Health, Bethesda, MD, USA, and were approved by the government of the Oberpfalz, Germany. In all *in vivo* experiments, the experimenter was blind to the treatment. Group sizes were estimated upon power analysis, based on results from previous publications (Jurek et al., 2012; Peters et al., 2014). Animals were randomly assigned to experimental groups, complying with equal mean body weight between groups.

Following surgery, the animals received a subcutaneous injection of antibiotics (0.03 ml enrofloxacin; 100 mg/1ml) and Buprenorphine as analgesic (0.05 mg/kg). Rats that received an acute infusion underwent surgery as described below, single-housed, and allowed to recover for at least 5 days. Animals were handled daily to habituate them to the infusion procedure in order to avoid non-specific stress responses during the experiment.

Locomotion was determined by Noldus EthoVision XT 14 software (distance travelled) and controlled by manual counting of line crossings.

### Chronic OXT infusion in male and female rats

To determine the effect of chronically infused OXT on parameters of interest (anxiety-like behavior, expression of OXT, OXTR, V1a, MEF2A, various stress-related genes (PCR array; see below) and MEF2A activity), osmotic minipumps (Alzet, model 1002, flow rate of 0.25 µl/hour for 14 days) filled with 40 µM or 4 µM OXT (equals 10 ng/h or 1 ng/h OXT) were implanted subcutaneously. The implant was placed in the abdominal region *via* a 1 cm long incision at the neck of the rat and connected *via* a silicone tubing with the i.c.v. cannula. The skin of the neck and the skull was closed with kallocryl. Rats were weighed on day 1, 8, and 13 of infusion, and 4 days after the infusion ended in the according experiment. After 13 days of chronic OXT infusion, rats were mildly stressed by elevated platform stress as described previously (Blume et al., 2008; Neumann et al., 2000a). The LDB was performed on day 14 of chronic OXT as previously described (Peters et al., 2014; Slattery and Neumann, 2010; Waldherr and Neumann, 2007). A second cohort of animals was tested at day 14 in the LDB without previous open arm exposure. For LDB testing, rats were placed in the light compartment and distance moved and percent time spent in each compartment during the 5-min test were assessed *via* a camera located above the box and analyzed by Noldus.

Acute i.c.v. VEH or OXT infusion were conducted as previously described (Jurek et al., 2015; Jurek et al., 2012).

### Social preference test

To assess social preference, rats were placed in an open field box (80 cm x 80 cm, 100 lux in the inner zone: 40 cm x 40 cm) and allowed to freely explore for 5 min. After habituation, an empty cage (non-social stimulus) was placed into the inner zone of the box for 5 min. The empty cage was then replaced by a cage with a conspecific (social stimulus) for another 5 min. Duration of direct contact (sniffing) was measured using JWatcher observational software (V 1.0, Macquarie University and UCLA) and compared to the investigation time of the non-social stimulus.

### Antisauvagine-30 and Stresscopin infusion

For acute local infusion of antisauvagine-30 (ASV) or stresscopin (SCP), 21-G guide cannulae were implanted stereotaxically 2 mm above both the left and right paraventricular nucleus (PVN) (−1.4 mm bregma, −1.8 or +2.1 mm lateral, 6.0 mm below surface of the skull; angle 10°; (Paxinos, 1998)) of the hypothalamus for subsequent infusion of ASV. The day before the experiment, rats were subjected to a mild stressor by placing them on an elevated platform for 5 min, to induce membrane incorporation of the CRFR2. On the day of experiment, male rats were injected with ASV (280 µM corresponding to 0.14 nmol/ 0.5 µl) bilateral into the PVN 10 minutes before the behavioral testing and returned to the homecage until testing in the LDB. SCP (1.4 mM, corresponding to 3 µg/ 0.5 µl per side) was injected 25 min prior to behavioral testing.

### Infusion of GapmeRs or TSBs for sCrfr2α knockdown or overexpression

The antisense LNA GapmeR and TSB were produced according to the custom design generated by a proprietary design software for optimal performance. 5’-FAM modification for transfection site control is indicated within the product sequence below. Both antisense oligonucleotides (ASO) contain phosphorothioate backbone modifications indicated by “*” in the product sequence.

Intra-PVN infusion of GapmeRs (0.5 nmol/ 0.5 µl, *in vivo* ready) targeted against the alternative splice variant of the CRFR2 was executed using the following coordinates (−1.7 mm bregma, ±0.3 mm lateral, 8.2 mm below surface of the skull). TSBs were infused similarly at the same molarity. The infusion site was closed with sutures and rats were allowed to recover for 7 days. Behavioral testing was executed 7 days after ASO infusion. Rats were subjected to a mild stressor paradigm, i.e. elevated platform for 5 min the day before behavioral testing in the LDB or social preference test.

Sequence of GapmeRs, TSBs and scrambled control are listed in the Key Resource Table.

### Tissue, plasma, and whole organ processing

After final behavioral testing, rats were immediately decapitated and the brains removed either for protein and RNA isolation from regions of interest or for freezing in methylbutane for later cryosectioning for histological verification of correct cannula placement or in situ hybridisation. Alternatively, after the behavioral test rats were anaesthetized with a lethal dose of urethane (2∼3 ml per rat), the cisterna magna punctuated, and CSF taken with a Hamilton syringe. After CSF was taken, rats were decapitated for trunk blood collection. Left and right adrenal glands, heart, and the thymus were removed, pruned from fat, and weighed. Oil-red staining of lipid vesicles in the adrenal cortex was conducted as described previously (Fuchsl et al., 2013). Trunk blood was collected, centrifuged for 10 min at 5000 rpm and 4 °C in EDTA-tubes, the supernatant was removed and kept as plasma in protein-low-binding tubes (Sarstedt, Nümbrecht) at −80°C for later ELISA analysis for ACTH and CORT levels.

### Cell lines and primary cultures

H32 cells are rat embryonic hypothalamic neurons that express the OXTR, V1a, MEF2A and C and the CRFR2, but not CRFR1. Authentication of cells was executed on the basis of OXTR sequencing, morphology, and controlled transcriptomics. Cells were cultured in DMEM/F12 with 10% heat inactivated FBS advanced, and Penicillin/Streptomycin. Cells were sub-cultured by gentle trypsination at 80% confluence. For transcriptional analyses, cells were seeded at a density of 3 ⨯ 10^6^ cells in a 25 cm^2^ cell culture dish the day before experiment, pre-incubated in serum-free stimulation medium for 1 h and stimulated with the respective treatment.

Primary neuronal and glial cultures were obtained from embryonic day 18 rat hypothalami, as described in detail in (Jurek et al., 2015).

### In silico analysis of MEF2 targets

Analysis of potential MEF2 targets by assessment of DNA binding regions was performed using the Geneious prime software (Geneious prime 2019.0.3; https://www.geneious.com).

### siRNA-mediated knockdown of MEF2A

H32 cells were seeded 24 h prior to transfections. For the MEF2A knockdown, both siRNA and scrRNA (Origene) were used at a concentration of 1 nM. Cells were transfected with Lipofectamine RNAiMAX Reagent (Invitrogen by Life Technologies) and incubated for 72 h at 37 °C and 5% CO_2_. After 48 h, the cells were stimulated with either 100 nM OXT or VEH for another 24 h.

### Chromatin -IP

Chromatin-IP was conducted using a MEF2A specific antibody. Cells were stimulated as described above, MEF2A-DNA complexes were fixated (3 % Glyoxal + 20 % Ethanol + 500 µl Acetic acid, pH 4.5) for 10 min, lysed, sheared by sonication, precleared with Sepharose beads, and MEF2A complexes isolated with a specific MEF2A antibody. DNA fragments were identified by qPCR with primers directed against CRFR2-specific MEF2A binding sequences (see Key Resource Table).

### Nano-Glo^®^ HiBiT Extracellular Detection System

An 11-amino-acid peptide tag called HiBiT was fused to the mCRFR2α using CRISPR-Cas9. The integration site of the HiBiT-containing Ultramer^®^ ssDNA donor sequence was directly upstream of Exon 6 of the CRFR2α gene. Sequences of guide RNAs used in HibiT-CRISPR-Cas9 tagging procedure is provided in the Key Resource Table. The donor sequence was delivered together with the RNP complex (Alt-R^®^ CRISPR-Cas9 system) by cationic lipid delivery using Lipofectamine™ RNAiMAX Transfection Reagent.

All further membrane-bound mCRFR2α was visualized using the Nano-Glo^®^ HiBiT Extracellular Detection System according to the manufacturer’s instructions. Luminescence was measured at the GloMax Explorer (Promega, Mannheim, Germany).

### Immunofluorescent labelling for co-localization of sCRFR2, OXTR, and OXT in OXTR-reporter mice

OTR-reporter mouse brains (OXTR-Venus, Yoshida et al., 2009) were kindly provided by Prof. Nishimori. For immunostaining, 40 µm thick cryo cut sections containing the PVN were permeabilized, blocked (PBS with 2 % BSA, 1 % Glycine, and 0.3 % TritonX-100, 1 h at room temperature), double labelled for sCRFR2α, in combination with OXT-Neurophysin I and Venus-OXTR or V1a/b (antibody details in table 2). The sections were gently transferred to SUPERFROST^®^ glass slides and mounted with Surgipath Premier coverslips using ProLong Glass Anti-Fade Reagent with NucStain (# P36984, Thermo Fisher Scientific). Images were acquired using a SP8 CLSM confocal microscope (Leica). Colocalization was analyzed by means of the BioVoxxel Version of Fiji using the plugin JACoP. Antibody specificity was assessed by pre-incubation with immunizing peptide, and in knockdown and overexpression systems (Fig S3).

### Immunocytochemistry (ICC) in primary hypothalamic rat neurons

0.05 ⨯ 10^6^ primary hypothalamic cells were seeded in BD Falcon Chamber slides (Poly-L-Lysine coated), fixated in 3 % Glyoxal, rinsed in PBS-T, and blocked with blocking solution (PBS, 2 % BSA, and 0.5 % Triton X-100, 1 % Glycine, 0.5 % cold water fish gelatine) for one hour followed by primary antibody (see table 1) incubations in PBS-T (0.1 % Triton X-100) overnight and secondary Alexa Fluor antibodies for 2 h in PBS-T.

### Western Blot and Dot Blot analysis

Western Blot and Dot Blot analysis was conducted as previously described (Jurek et al., 2015; Martinetz et al., 2019; Meyer et al., 2018). Briefly, 25 µg of whole cell extract were loaded and separated in a 10 % Mini PROTEAN or Criterion TGX Stainfree gel (BioRad). Western blot analysis was used to detect total protein and phosphorylated forms of various proteins (see Key Resource Table), using the Stainfree total protein method (BioRad, Steinheim, Germany) as loading control.

For Dot Blot analysis, 10µg of total protein was pipetted onto a Nitrocellulose membrane, allowed to dry and processed identical to the Western Blot protocol. Loading was controlled by Ponceau red staining.

### Radiolabeled *in situ* hybridization for OXT mRNA content in the PVN

To assess hypothalamic OXT mRNA content in rats treated chronically with OXT, *in situ* hybridization was conducted as described previously for mice (Peters et al., 2014). The sequence of the ^35^S-labeled probe (provided in the Key Resource Table) is specific for rats and mice. Briefly, 16µm cryo-sections were mounted onto pre-coated SuperFrost Plus Slides (Menzel-Gläser, Braunschweig) and processed as described previously (Peters et al., 2014). Brain slices, which contained comparable sections of PVN, were measured for each subject to provide individual means. Expression of OXT mRNA was measured as grey density with ImageJ 1.51 g. Background activity was automatically subtracted from measured areas to yield values for specific binding.

### Receptorautoradiography (RAR) for OXTR and V1aR binding

To measure whether chronic OXT altered hypothalamic V1aR and or OXTR binding, rats were decapitated, brains were removed, quickly frozen in pre-chilled n-methylbutane on dry ice, and stored at −20 °C. Brains were cut into 16-mm coronal cryostat sections and mounted on slides. The receptor autoradiography procedure was performed according to Lim et. al. (Lim et al., 2004) using a linear V1aR antagonist ^125^I-phenylacetyl-d-Tyr(Me)-Phe-Gln-Asn-Arg-Pro-Arg-Tyr-NH2 (Perkin Elmer, USA) or a linear OXTR antagonist [^125^I]-d(CH_2_)^5^[Tyr(Me)^2^-Tyr-Nh_2_]^9^-OVT (Perkin Elmer, USA) as tracers. The optical density of V1aR and OXTR was measured using ImageJ (V1.51g). Receptor density was calculated per rat by taking the mean of bilateral measurements of four to six brain sections per region of interest. After subtraction of tissue background, the data was converted to dpm/mg (desintegrated points per minute/milligram tissue) using a [125I] standard microscale (Amersham, Germany).

### ACTH and CORT ELISA

Approximately 1 ml of trunk blood was collected in EDTA-coated tubes on ice (Sarstedt, Nümbrecht, Germany) and centrifuged at 4 °C (5000 rpm, 10 min). Supernatant was removed and stored at −80°C as plasma samples, until assayed using a commercially available ELISA for ACTH (analytical sensitivity 0.22 pg/ml, intra-assay and interassay coefficients of variation ≤ 7.1 %, IBL, Hamburg) and CORT (sensitivity <1.63 nmol/l, intra-assay and inter-assay coefficients of variation ≤ 6.35 % IBL, Hamburg) and a 96 well plate reader (Optima FluoStar, BMG) (33).

### RNA isolation for qPCR and PCR array

To analyze mRNA expression of target genes in specific brain regions, rat tissue (PVN, hippocampus, prefrontal cortex) was punched and frozen in 1 ml TriFast Gold (PeqLab) at −80 °C until further processing. For RNA isolation the tissue was thawed on ice, homogenized using a pestle (VWR, Radnor, USA), and RNA was isolated according to the protocol provided by the manufacturer with some modifications as described previously (Jurek et al., 2015).

To isolate RNA from the stimulated cells, the medium was aspirated, cells were washed with PBS, and RNA was isolated according to the instruction manual of the Protein/RNA isolation Kit by Macherey Nagel.

300 ng of total RNA per sample were used for reverse transcription into cDNA using Super Script IV First strand Synthesis System for RT-PCR (Invitrogen). Relative quantification of MEF2A, MEF2C transcript variant X1, sCRFR2α, and CRFR2α mRNA levels was performed using SYBR Green (QuantiFast Qiagen), using ribosomal protein L13A (Rpl13A,) and cyclophilin A (CycA,) as housekeeping genes (Bonefeld et al., 2008). Primer efficiency for each primer pair was calculated by serial dilution of test cDNA using the Pfaffl method (Bustin et al., 2009; Pfaffl, 2001). Specificity of the qPCR was assured by omitting reverse transcription and by using ddH_2_O as template. As the results obtained using Rpl13A and CycA yielded similar results, only those for Rpl13A are shown. The PCR protocol consisted of an initial denaturation step of 5 min at 95 °C, followed by 50 cycles of denaturation at 95 °C for 10 s, and annealing/extension at 60 °C for 45 s. At the end of the protocol, a melting curve was generated, and PCR products were analyzed by agarose gel electrophoresis to confirm the specificity of the primers. All samples were run in triplicate. The custom RT^2^ PCR array (330171 CLAR25389) was purchased from Qiagen and pipetted according to the manufacturer’s protocol.

### TransAM MEF2 binding kit

Protein samples were collected according to the manufacturer’s protocol (Active Motif, Rixensart), protein concentration was determined using a BCA assay, and 20 µg protein was loaded onto the pre-coated 96 well plate. The included antibody (panMEF2 subform unspecific) was replaced by MEF2A and MEF2C subform specific antibodies from OriGene/Acris (Key Resource Table). Fluorescence was determined after 10-15 min developing at 450 nm in a plate reader (FluoStar Optima, BMG)

### Quantification and statistical analysis

Parametric one-way (factor treatment) or two-way (factors treatment x time) analysis of variance (ANOVA), followed by the Holm Sidak post hoc correction, were performed for statistical analyses of behavioral and molecular experiments. Non-parametric data was analyzed by the Kruskal-Wallis ANOVA on ranks and the Tukey post hoc test. Separate parametric t-test between two groups or non-parametric Mann-Whitney U tests (MWU) were performed. Statistical significance was accepted at p < 0.05. In behavioral experiments, “n” represents the number of animals. Data are presented as mean ± or + standard error of the mean (SEM), as indicated in the figure legend. Statistical analyses were performed using Sigma Plot (version 11.0.0.75, Systat Software).

### Key resources table

## AUTHOR CONTRIBUTIONS

Conceptualization, B.J., S.P., D.A.S., E.H.vdB., I.D.N.; Methodology, B.J., J.W., M.M., I.B.; Validation, J.W., M.M., I.B., K.K., M.B., M.R., A.K.S., A.B.; Investigation, J.W., M.M., I.B., S.P., M.R., D.L., K.K., K.H.

O.J.B., M.B.; Writing – Original Draft, J.W., B.J.; Writing – Review & Editing, S.P., S.O.R., O.J.B., D.A.S, E.H.vdB., I.D.N., B.J.; Funding Acquisition, I.D.N., B.J.; Resources, I.D.N; Supervision, B.J., S.P., E.H.vdB., I.D.N.

## ACKNOWLEDGEMENTS

We would like to thank Joan Vaughan and Prof. Dr. Harold Gainer for providing sCRFR2 and OXT-Neurophysin I antibodies, Rodrigue Maloumby, Vinicius Oliveira, and Andrea Havasi for excellent technical help, and Carl-Philipp Meinung and Serena Gusmerini for excellent scientific data. This work was supported by the German Research Foundation GRK 2147 (IDN, BJ, OJB), JU3039/1-1 (BJ), NE 465/27-1 (IDN), NE 465/31-1 (IDN), NE 465/19-1 (IDN, EvdB), and EU (FemNat-CD; IDN).

The funders had no role in study design, data collection and analysis, decision to publish, or preparation of the paper.

The authors declare no competing interests.

## SUPPLEMENTARY

**Table S1.**
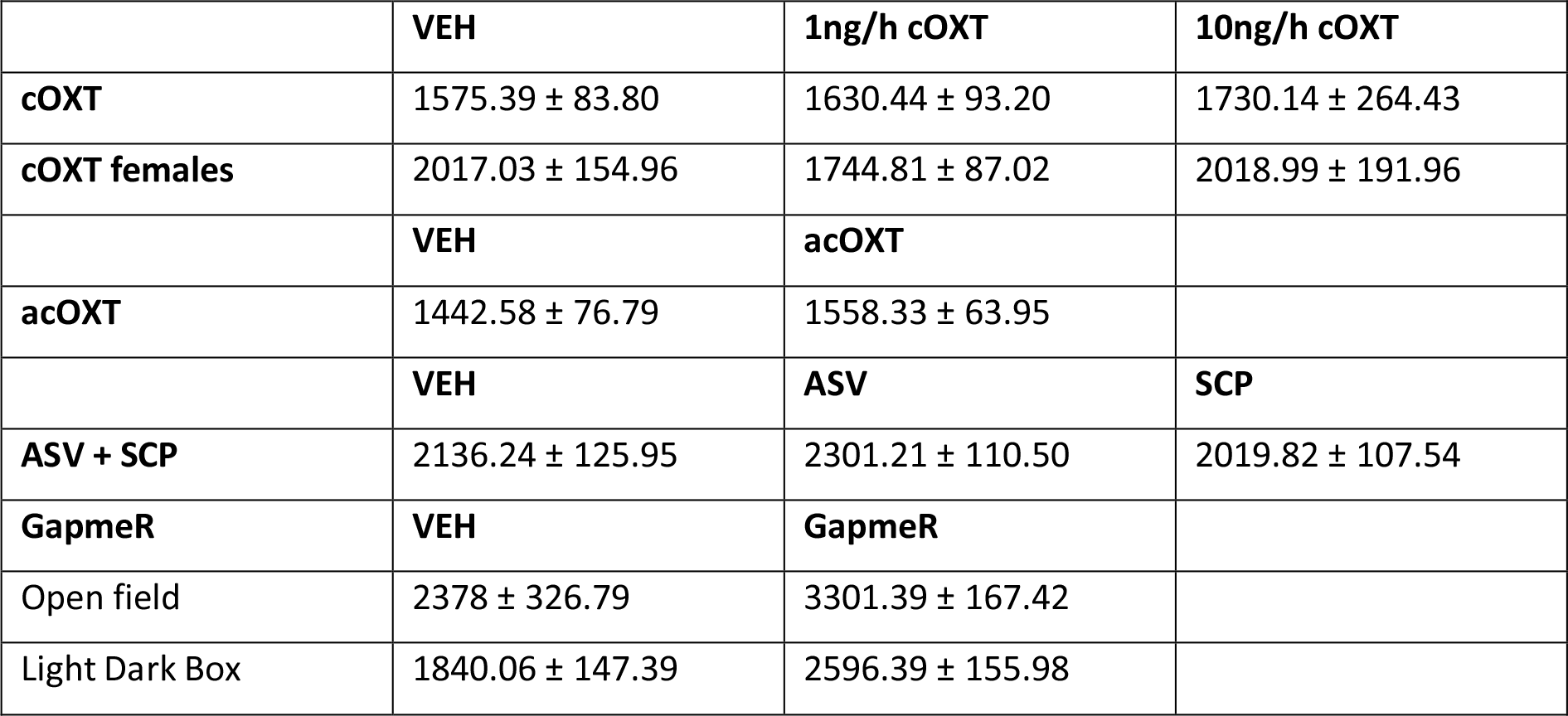
Locomotion assessed in behavioral tests in [cm] by Noldus EthoVision.

**Table S2.**
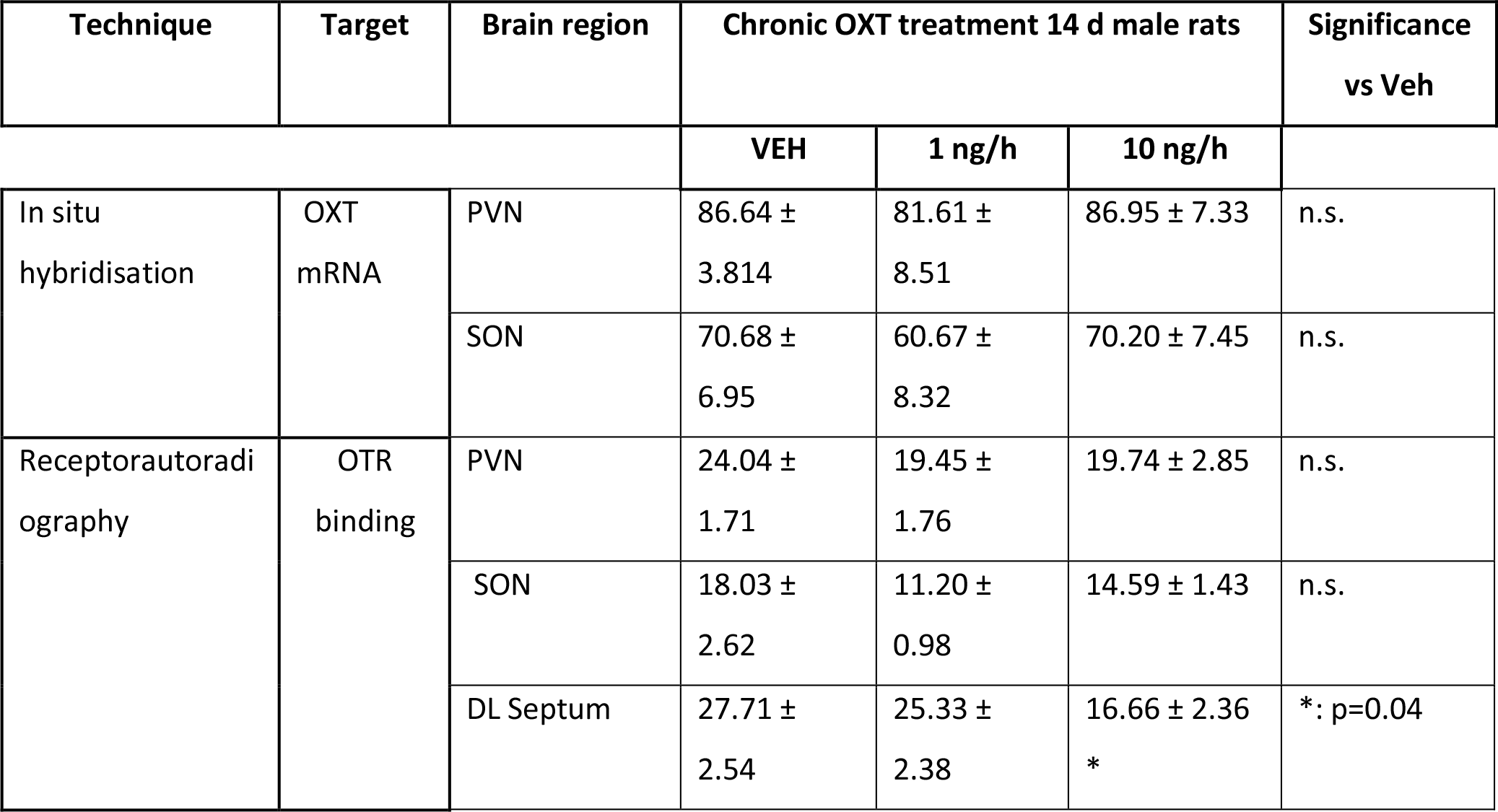
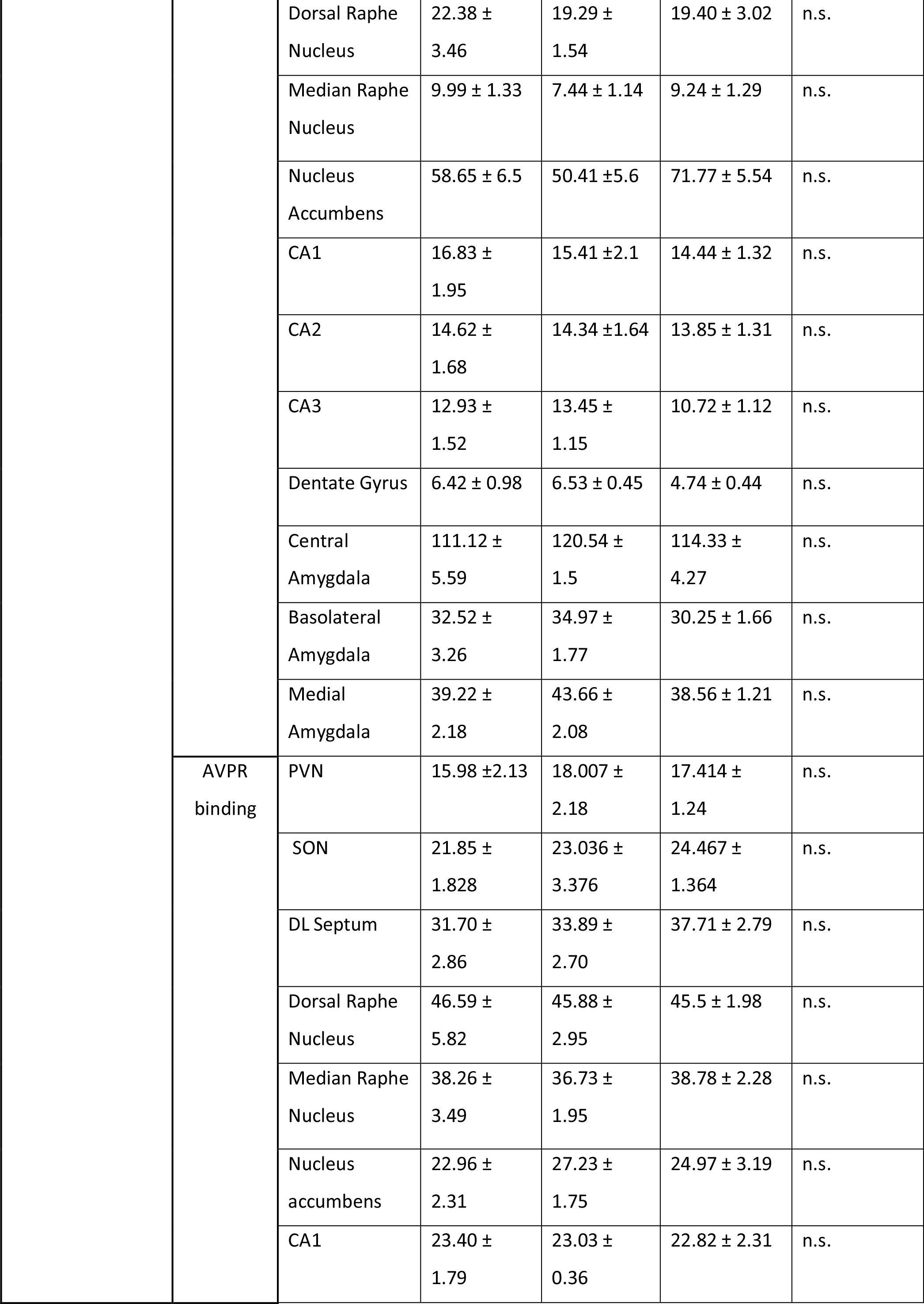
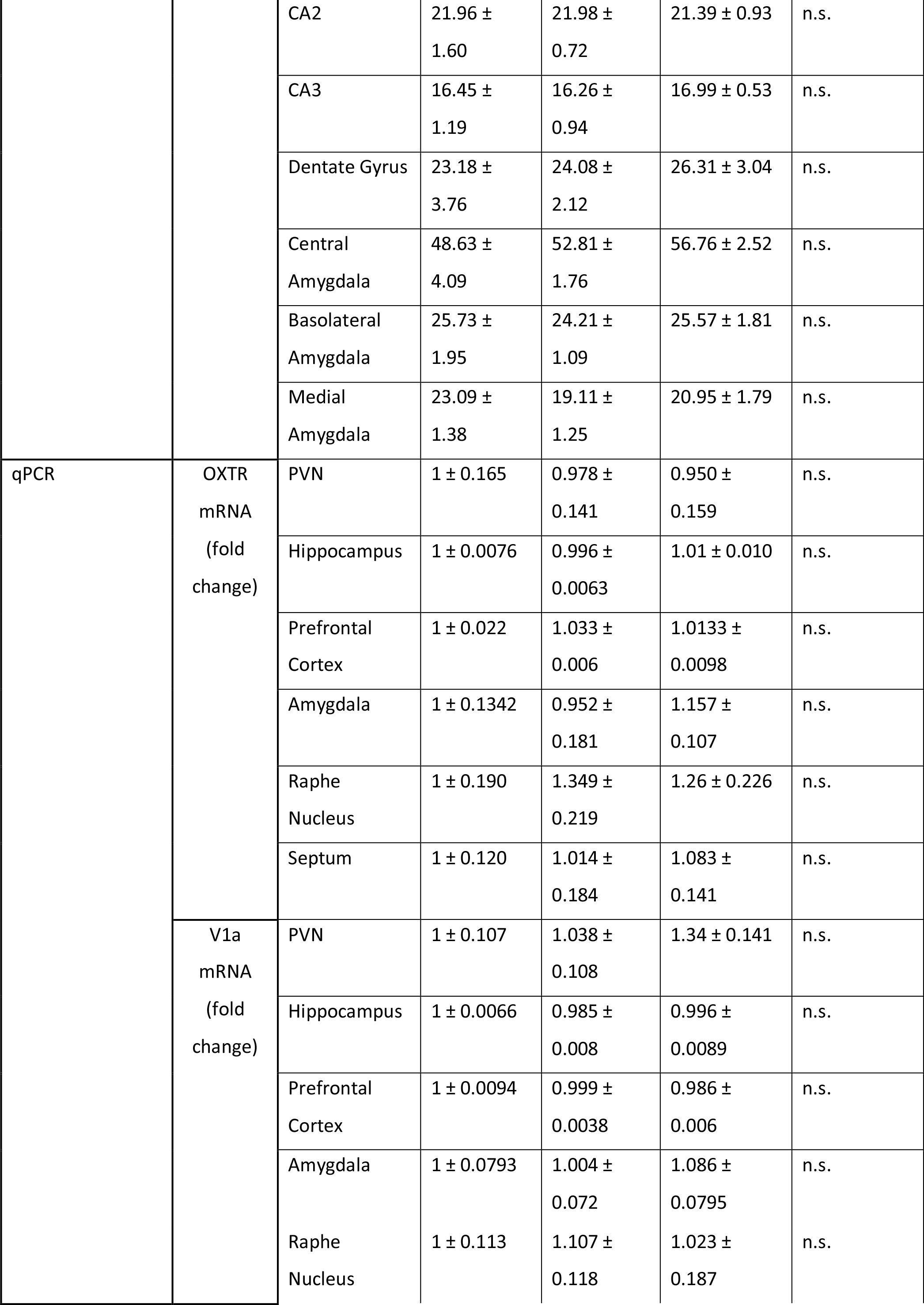
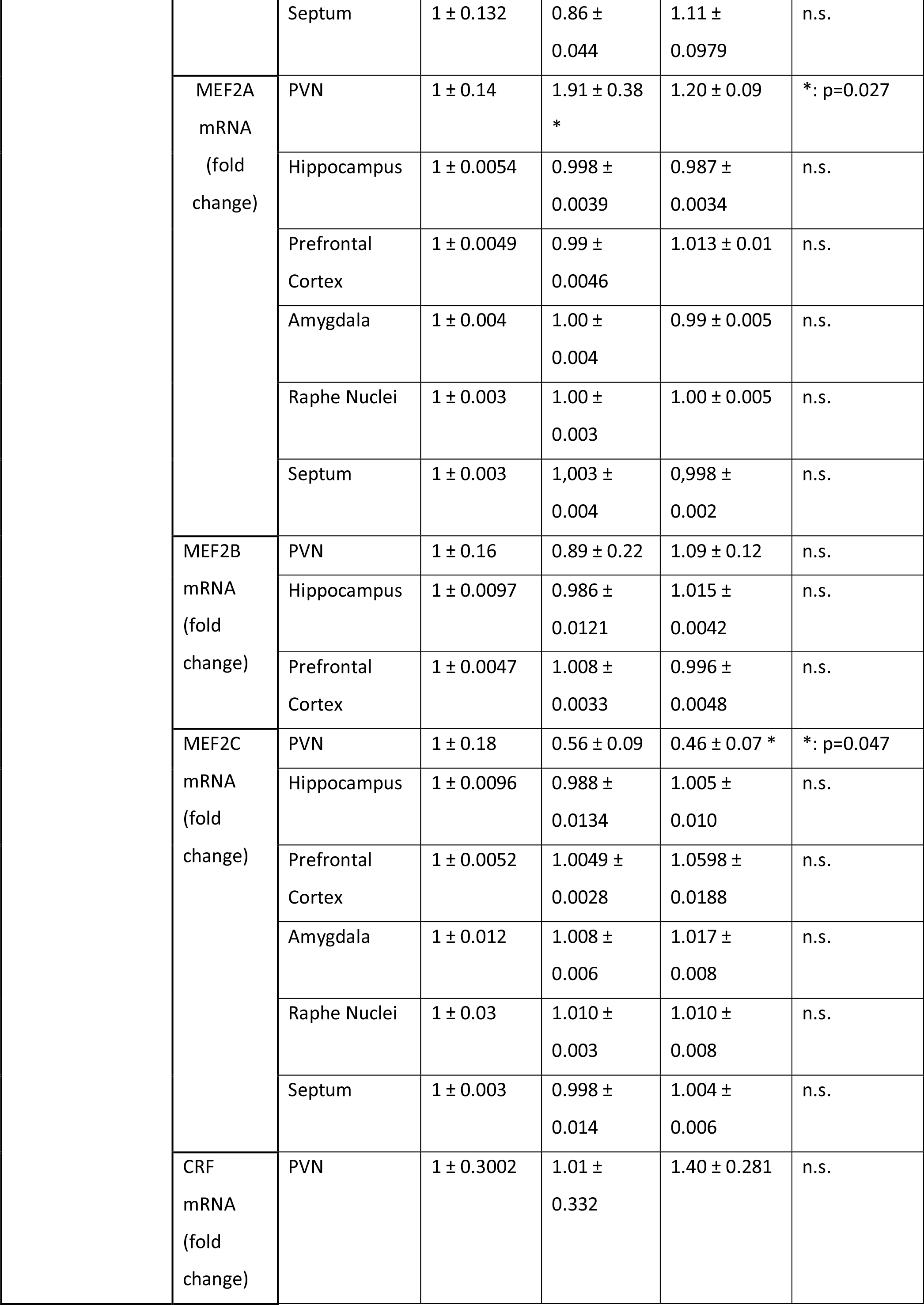
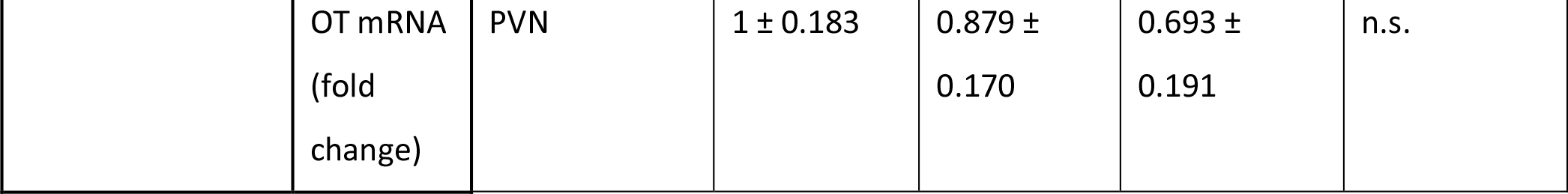
Central effects of chronic OXT on gene or protein expression in stress- and anxiety-related brain regions evaluated by in situ hybridization, receptorautoradiography, or qPCR. N.s.= not significant, n= 6-10 male rats, depending on the technique and experiment.

**Table S3.** Influence of chronic OXT treatment on peripheral parameters. None of the parameters assessed differed significantly between treatment groups.

|  | VEH | 1 ng/h chronic OXT | 10 ng/h chronic OXT |
| --- | --- | --- | --- |
| plasma ACTH [pg/ml] | $53.056 \pm 11.510$ | $64.67 \pm 14.41$ | $50.03 \pm 11.69$ |
| plasma Cort with EPF but no LDB [ng/ml] | $389.29 \pm 45.03$ | $393.11 \pm 58.06$ | $463.53 \pm 46.92$ |
| Plasma Cort with EPF and with LDB [ng/ml] | $935.48 \pm 59.84$ | $638.53 \pm 131.44$ | $716.11 \pm 139.65$ |
| Plasma Cort no EPF but with LDB [ng/ml] | $839.54 \pm 67.76$ | $594.01 \pm 74.00$ | $805.99 \pm 89.63$ |
| rel. adrenal weight [mg/g body weight] | $0.192 \pm 0.015$ | $0.179 \pm 0.012$ | $0.199 \pm 0.014$ |
| absolute adrenal weight left [g] | $0.036 \pm 0.003$ | $0.0354 \pm 0.0013$ | $0.0377 \pm 0.0022$ |
| absolute adrenal weight right [g] | $0.0342 \pm 0.0022$ | $0.0305 \pm 0.0017$ | $0.0352 \pm 0.0016$ |
| adrenal cortex lipid droplet content (oilred staining) [%] | $8.9 \pm 0.01$ | $8.9 \pm 0.01$ | $6.85 \pm 0.01$ |
| heart weight [g] | $0.9608 \pm 0.0155$ | $1.0965 \pm 0.0401$ | $1.0818 \pm 0.0151$ |
| body weight gain [g] | $14.4 \pm 1.5$ | $16.8 \pm 0.80$ | $16.3 \pm 0.42$ |
| rel. thymus weight [mg/g] | $1.38 \pm 0.10$ | $1.22 \pm 0.06$ | $1.32 \pm 0.081$ |
| Dermal OXTR expression (fold-change mRNA) | $1.000 \pm 0.03$ | $0.952 \pm 0.015$ | $0.923 \pm 0.038$ |

**Figure S1.**
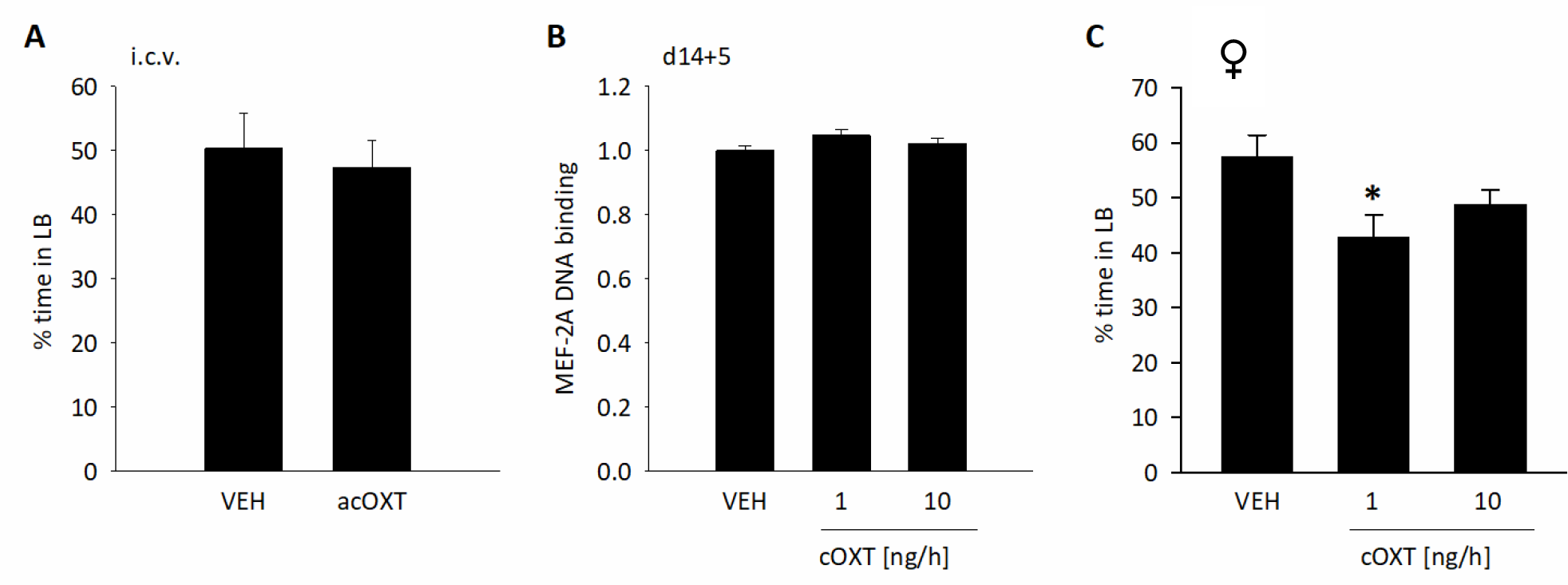
Continuative behavioral data. (A) Anxiety-like behavior of male rats in the LDB 10 min after i.c.v. OXT infusion at a dose of 0.1µg/5µL. Data are represented as mean % time spent in the LB + SEM. t = 0.420, one-tailed p-value = 0.339; n_(VEH)_ = 14, n_(OXT)_ = 14. (B) When tested for anxiety-like behavior in the LDB 5 days after the cOXT infusion ended, % time spent in the LB returned to basal levels in the treatment groups. Data are represented as mean % time spent in the LB + SEM. n_(VEH)_ = 5, n_(1ng cOXT)_ = 5, n_(10ng cOXT)_ = 5. (C) Anxiety-like behavior of female rats increases in the LDB after 14 days of i.c.v. of chronic OXT at a dose of 1ng/h, but not 10ng/h. Data are represented as mean % time spent in the LB. Kruskal-Wallis, H = 7.525, * p = 0.023 vs VEH, n_(VEH)_ = 6, n_(1ng cOXT)_ =7, n_(10ng cOXT)_ = 7.

**Figure S2.**
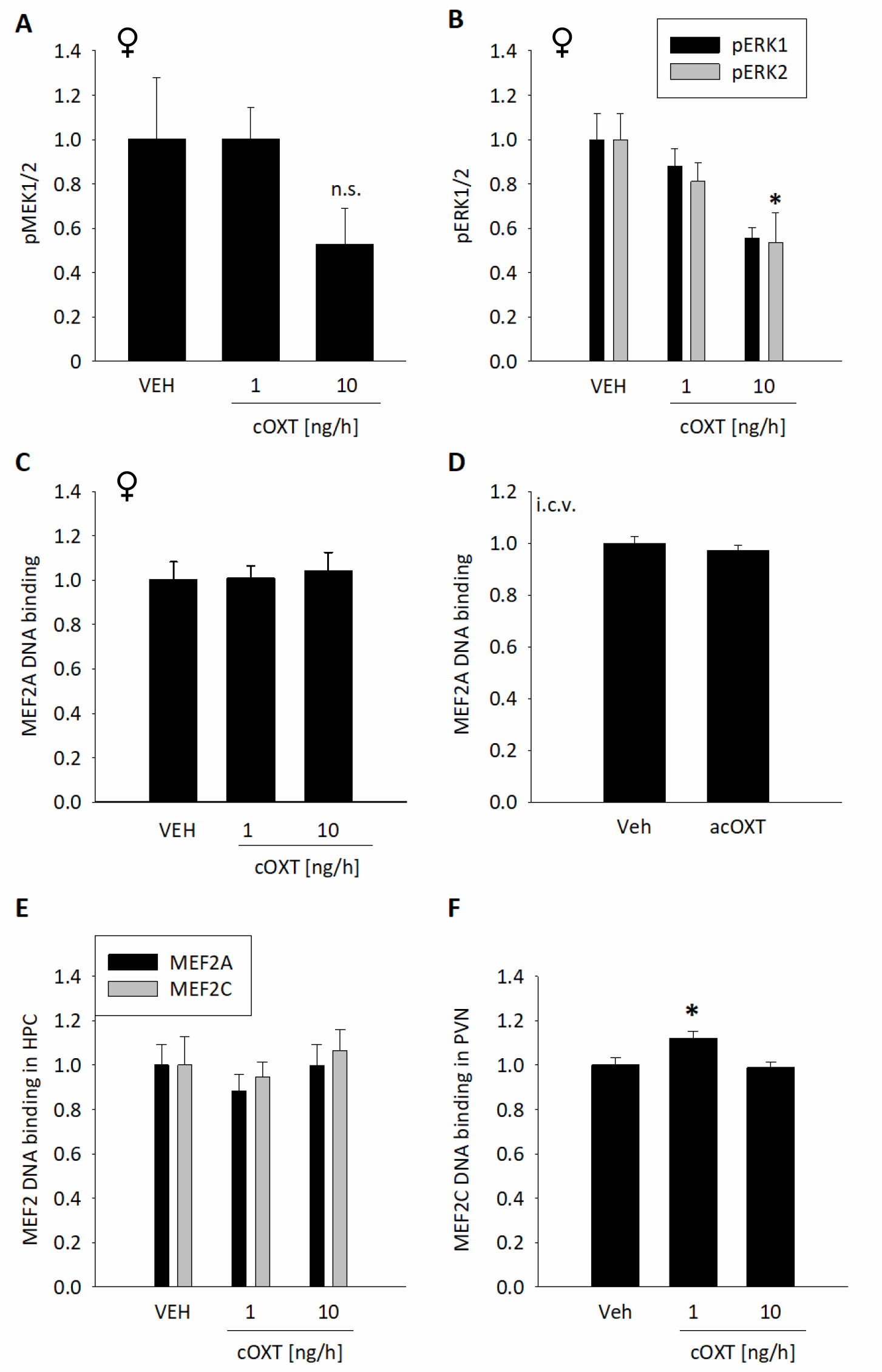
Intracellular signaling in the PVN or hippocampus of male and female rats. (A) MEK1/2 phosphorylation in the PVN of female rats tended to decrease in the 10ng/h cOXT group without reaching significance. Data are represented as fold changes vs VEH + SEM. n_(VEH)_ = 6, n_(1ng cOXT)_ =7, n_(10ng cOXT)_ = 7. (B) Phosphorylation of ERK2 in the PVN of female rats decreases significantly in the 10ng/h cOXT group, whereas ERK1 tends to decrease. No effect on ERK1/2 phosphorylation was observed in the 1ng/h cOT group. Data shown as fold changes vs VEH + SEM. pERK1: F_(2,16)_ = 3.293, p = 0.067; pERK2: H = 5.795, * p = 0.045; n_(VEH)_ = 6, n_(1ng cOXT)_ =7, n_(10ng cOXT)_ = 7. (C) No significant effect of cOXT treatment on MEF2A DNA-binding in the PVN of virgin female rats has been found. Data are represented as fold changes in DNA binding activity + SEM F_(2;22)_ = 0.177; p = 0.839; n_(VEH/1 ng/h cOXT)_ = 8, n_(10 ng/h cOXT)_ = 7. (D) No effects of acute i.c.v. OXT infusions have been observed on MEF2A binding activity in the PVN of male rats. Data are represented as mean fold changes (+SEM) of DNA binding activity compared to VEH. Mann-Whitney U = 8.000, p = 0.247; n_(VEH)_ = 5, n_(OXT)_ = 6. (E) No significant changes in MEF2A and MEF2C binding activity in the hippocampus of male rats have been detected after chronic OXT treatment. Data are represented as fold changes in DNA binding activity + SEM, indicated by binding of MEF2A *in vitro* to its responsive element and fluorescent antibody-labeling of bound MEF2A. MEF2A: F_(20,20)_ = 0.632, p = 0.543, all treatment groups = 7. MEF2C: F_(20,20)_ = 0.347, p = 0.712. (F) In contrast to MEF2A (Fig. 2E), MEF2C binding activity increased slightly by 1ng/h, but not 10ng/h cOXT. Data are represented as fold changes in DNA-binding activity + SEM in the PVN of male rats. F_(2;17)_ = 5.633, p = 0.015, Holm Sidak *p = 0.015 vs VEH; n_(VEH)_ = 6, n_(1ng cOXT)_ = 6, n_(10ng cOXT)_ = 6.

**Figure S3.**
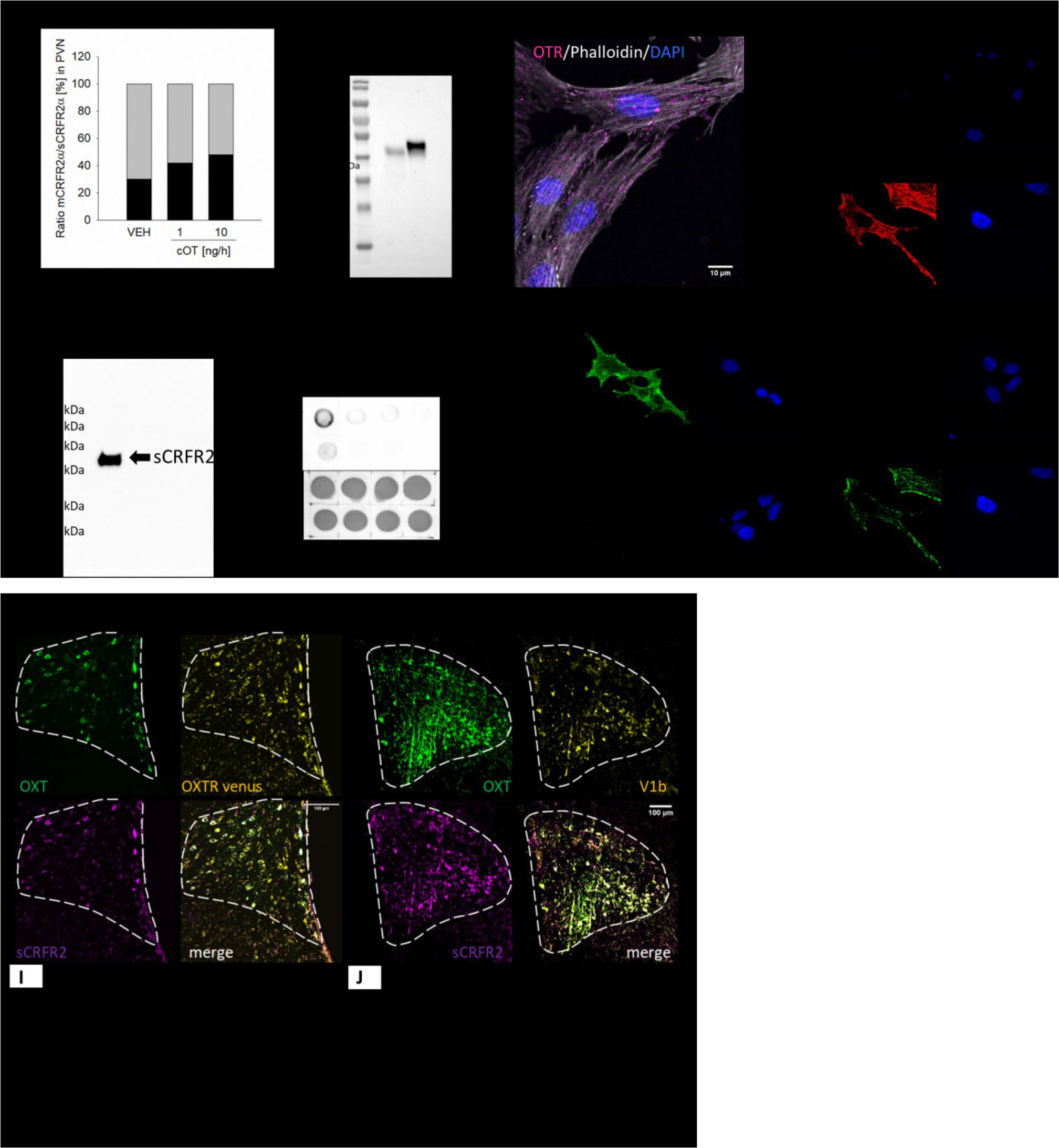
Antibody specificity and OXTR expression in H32 cells and *in vivo* co-localization analyses. (A) Protein ration of mCRFR2α over sCRFR2α expression in PVN tissue lysates of virgin female rats treated with 1 or 10 ng/h chronic OXT for 14 days. Unlike in male PVN tissue, no significant differences were detected. (B) Western Blot showing the input control, MEF2A precipitated lysate and antibody-Isotype-control precipitation of the chromatin immunoprecipitation from H32 cells as described in the results section. One single band in the second lane indicates a successful pull-down of MEF2A with cross-linked DNA, which has been analyzed by qPCR using primers covering MEF2A binding sites in Exon 2 and Exon 6. Antibody-Isotype pull down indicates specificity of the MEF2A antibody and is used for background signal subtraction. “Input” serves as loading control for the enrichment by the MEF2A antibody. (C) OXT receptor (OXTR) expression in H32 cells, OXTR in magenta, actin in white, DAPI in blue. Specificity of the Anti-OXT Receptor Antibody (AVR-013) from Alomone labs (Jerusalem, Israel) was tested and published elsewhere (Xiao et al., 2017). (D) OXTR staining in H32 cells treated with OXTR siRNA (OriGene Oxtr rat siRNA Oligo Duplex (Locus ID 25342)) in comparison to H32 wildtype cells. (E) To confirm specificity of the sCRFR2α antibody, we tested it in a Western Blot using rat plasma under standard denaturing conditions (SDS, 95°C for 5 min). The antibody produced one single band at around 38 kDa, which is the predicted molecular weight. The left panel shows a dot blot with native rat plasma samples and increasing amounts of blocking peptide. Ponceau red staining prior to antibody labeling served as loading control, and confirms even protein loading onto the membrane. (F) left: V1bR staining in H32 cells treated with a V1bR expression plasmid (OriGene, Avpr1b (Myc-DDK-tagged ORF) in comparison to H32 wildtype cells with no endogenous V1bR expression. right: V1aR staining in H32 cells treated with V1aR siRNA (OriGene, Avpr1b (Rat) −3 unique 27mer siRNA duplexes) in comparison to endogenous expression in wildtype cells. Scale bar represents 10 µm. (G) Representative stainings showing co-localization of OXT-neurophysin I (green), sCRFR2α (magenta), and OXTR-Venus (yellow) in the PVN (indicated by white dotted line) of male OXTR-Venus reporter mice. (H) Representative staining showing colocalization of OXT (green), sCRFR2α (magenta), and V1bR (yellow) in the PVN of male Wistar rats. (I) Pearson and Manders overlap coefficient of OXTR, OXT, and sCRFR2α positive neurons reveal substantial (45-64%) overlap between the OXTR, OXT, and sCRFR2α. (J) Pearson and Manders coefficient reveal partial (24-37%) overlap of V1a/b and sCRFR2α in the PVN. OXT-neurophysin and sCRFR2α show similar co-expression (∼60%) in rats and mice, indicating a comparable expression pattern of the sCRFR2α in both species.

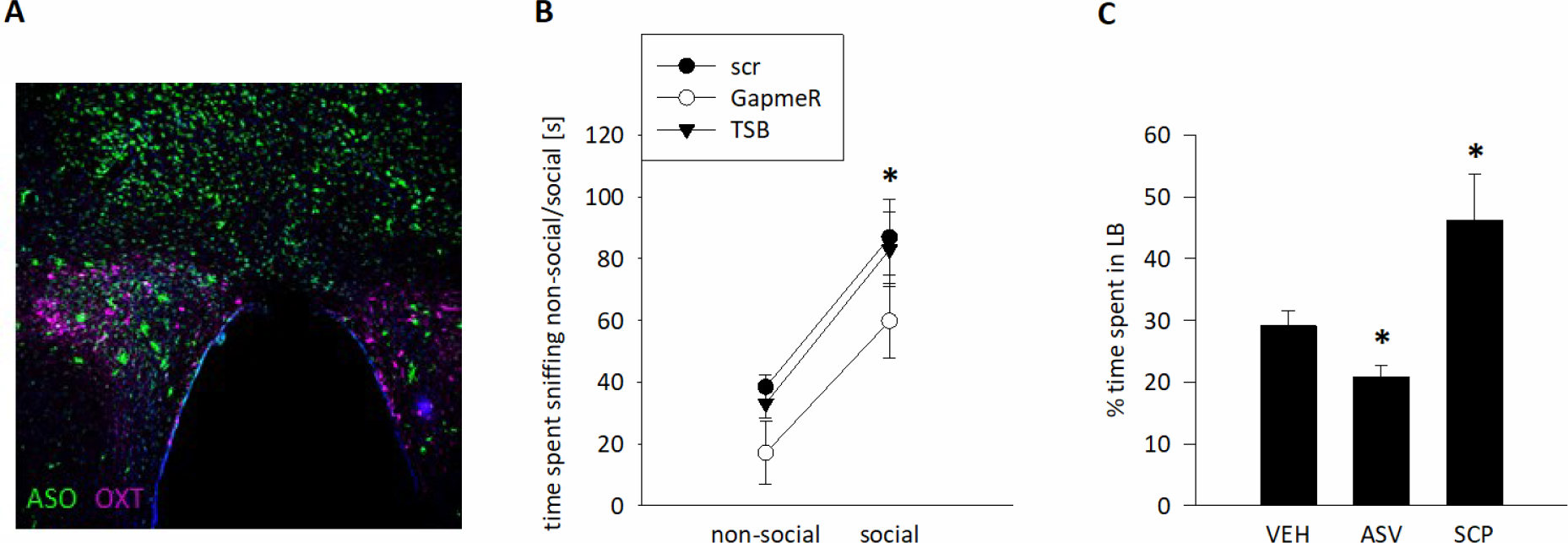
(A) Distribution of 5’-FAM-labelled antisense Oligos (ASOs) in the hypothalamic area after microinfusions (1nmol/0.5µl per animal). Direct infusions into the PVN (Bregma −1.7 AP, +/-0.3 ML, −8.2 DV) result in mostly PVN and dorsal hypothalamus staining. Scale bar represents 100µm, Magenta: OXT-Neurophysin positive neurons, green: GapmeRs. (B) Social preference measured by percentage of time spent investigating (sniffing) a non-social stimulus (empty cage) versus a social stimulus (cage with a conspecific) for 5 min each. All animals display social preference (scr, p = 0.001, GapmeR, p = 0.033, TSB, p < 0.01; Bonferroni’s post hoc analysis) with no significant effect of treatment. This indicates that the effect of the sCRFR2α on anxiety-like behavior has no social component. (C) Assessment of the anxiolytic effect of the PVN CRFR2α. Anxiety-like behavior of male rats increased in the LDB 10 min after intra-PVN administration of the CRFR2α-specific antagonist antisauvagine-30 (ASV; 280 µM), and decreased following infusion of the CRFR2α-specific agonist stresscopin (SCP; 1.4 mM). Data are represented as mean + SEM. F_(2;51)_ = 12.099; p < 0.001; Holm-Sidak * p < 0.03 vs VEH. n_(VEH)_ = 23, n_(ASV)_ = 21, and n_(SCP)_ = 8.

